# OncoPubMiner: A platform for oncology publication mining

**DOI:** 10.1101/2022.03.11.483968

**Authors:** Quan Xu, Yueyue Liu, Dawei Sun, Jifang Hu, Xiaohong Duan, Niuben Song, Jiale Zhou, Junyan Su, Siyao Liu, Fan Chen, Zhongjia Guo, Hexiang Li, Qiming Zhou, Beifang Niu

## Abstract

Knowledge bases that are up-to-date and of expert quality are fundamental in biomedical research fields. A knowledge base established with human participation and subjected to multiple inspections is crucial for supporting clinical decision-making, especially in the exponentially growing field of precision oncology. The number of original publications in the field has skyrocketed with the advancement of technology and in-depth research evolved. It has become an increasingly pressing issue that researchers need to consider how to gather and mine these articles accurately and efficiently. In this paper, we present OncoPubMiner (https://oncopubminer.chosenmedinfo.com), a free and powerful system that combines text mining, data structure customization, publication search with online reading, project-centered and team-based data collection to realize a one-stop “keyword in, knowledge out” oncology publication mining platform. It was built by integrating all the open-access abstracts from PubMed and full-text articles from PubMed Central, and is updated on a daily basis. The system makes it straightforward to obtain precision oncology knowledge from scientific articles. OncoPubMiner will assist researchers in developing professional structured knowledge base systems efficiently, and bringing the oncology community closer to achieving precision oncology goals.

**Graphical Abstract:** OncoPubMiner’s one-stop “keyword in, knowledge out” workflow (A) is built on key features such as text mining (B), publication search (C), form customization (D), and team-based curation (E).

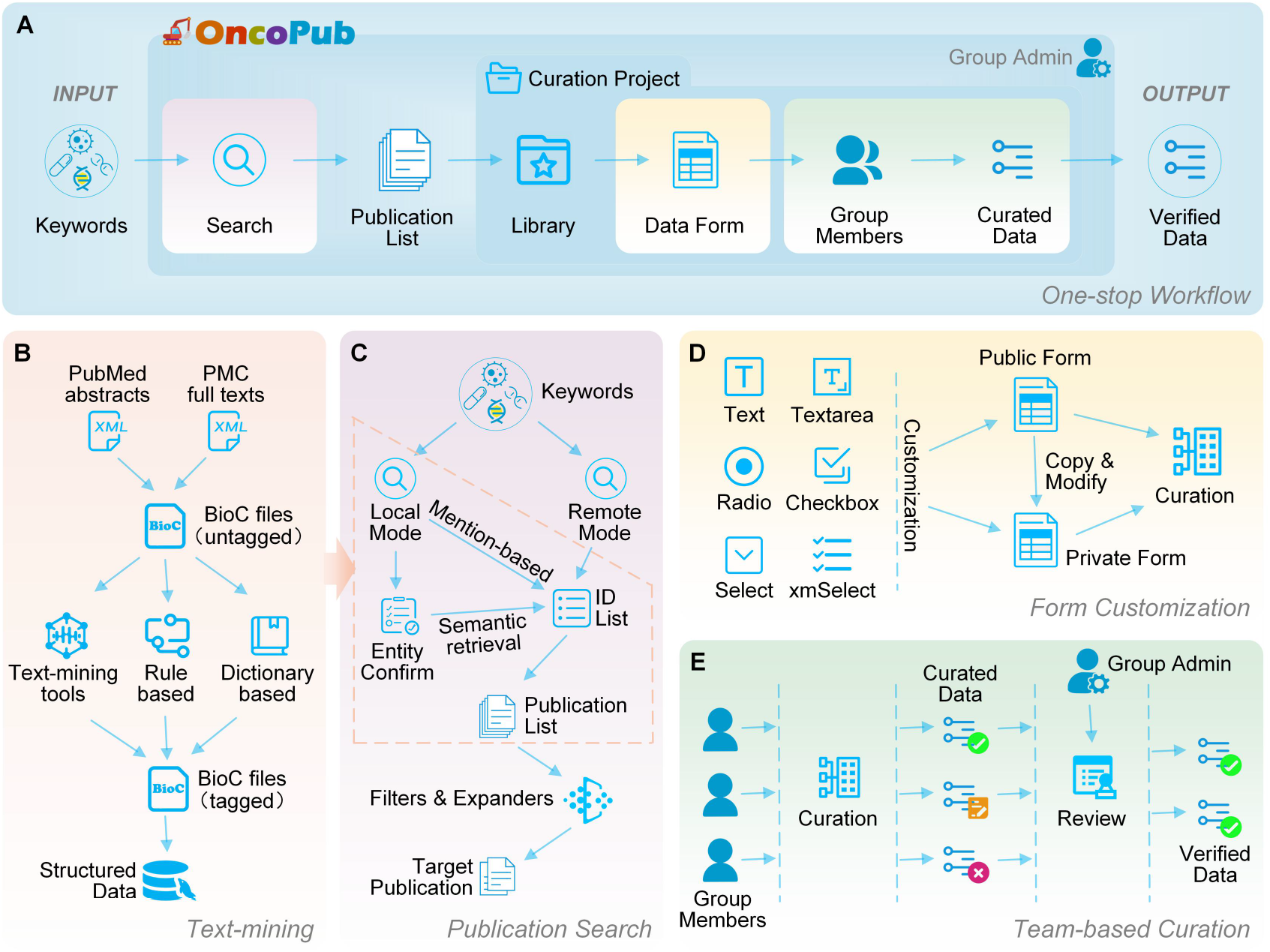

## INTRODUCTION

Studies on precision medicine in cancer have surged in recent years. Precision oncology knowledge bases built on top of the significant findings from these studies are critical for both oncologists and patients. They are the gold standard for developing new methodologies, tools, and algorithms that can assist new discoveries and clinical applications (1–6). Given the significance of knowledge bases in supporting cancer clinical decision-making, a large number of knowledge bases have been constructed, including Memorial Sloan Kettering’s OncoKB database, which was designated as the first tumor mutation database by the United States Food and Drug Administration (4,7–12). However, these established knowledge bases have some limitations such as not enabling free access, not allowing batch download, not providing continuous or real-time data updates, and the inability to be effectively utilized due to inappropriate data structure. Because of these constraints, many institutions have to establish their knowledge bases by curating original articles from scratch. As previously mentioned, the number of scholarly publications in precision oncology has increased tremendously. However, according to one study, more than 70% of researchers have tried but failed to repeat another scientist’s work, and more than half failed to replicate their own experiments (13). Therefore, researchers need to investigate multiple independent studies (14). Because literature retrieval and data mining are challenging (15), developing a database is time-consuming and labor-intensive (16–18). Many institutions are deeply involved in such process, considerably impeding the achievement of precision oncology goals.

Researchers already take some efforts in developing natural language processing algorithms and tools (19–24), as well as optimizing article retrieval (25–34). Most of these works are based on automatic text mining or literature retrieval methods. Textpresso Central and TeamTat are two platforms that recognize the necessity of team-based manual data mining (33,35). The solutions listed above can assist with data mining and knowledge base creation somewhat, yet they are away from adequate. Besides entity tagging and publication retrieval, effective article screening is also critical. Furthermore, a capacity that dynamically create data collection forms, as well as a platform that facilitates team collaboration in reading publications and collecting data, are expected to fulfill the fluctuating needs of different institutions for knowledge architectures.

In this paper, we present OncoPubMiner, a free platform for mining oncology publications. This one-stop “keyword in, knowledge out” (KI-KO) data collection platform combines text mining, data structure customization, article search with online reading, project-centered and team-based data collection. We downloaded open-access PubMed abstracts and PubMed Central (PMC) full-text articles, and then used scripts to monitor daily data updates. Natural language processing (NLP) technologies and a self-organized precision oncology vocabulary are employed to tag and standardize the entities. OncoPubMiner has a publication search engine for accurate literature retrieval based on semantics and mentions. It also provides a collaborative environment for the researcher teams to manage and curate publications and construct knowledge bases on precision medicine.

## SYSTEM DESCRIPTION

OncoPubMiner is a new, all-in-one publication mining platform based on KI-KO methodology. It analyzes open access publications from PubMed and PMC. It provides a publication search engine and facilitates data model customization. The curation project serves as a hub for a team of researchers to search, manage and read publications, to collect data, and to retrieve, review, and refine knowledge from the publication, in a one-stop KI-KO workflow (Fig. 1).

**Figure 1.**
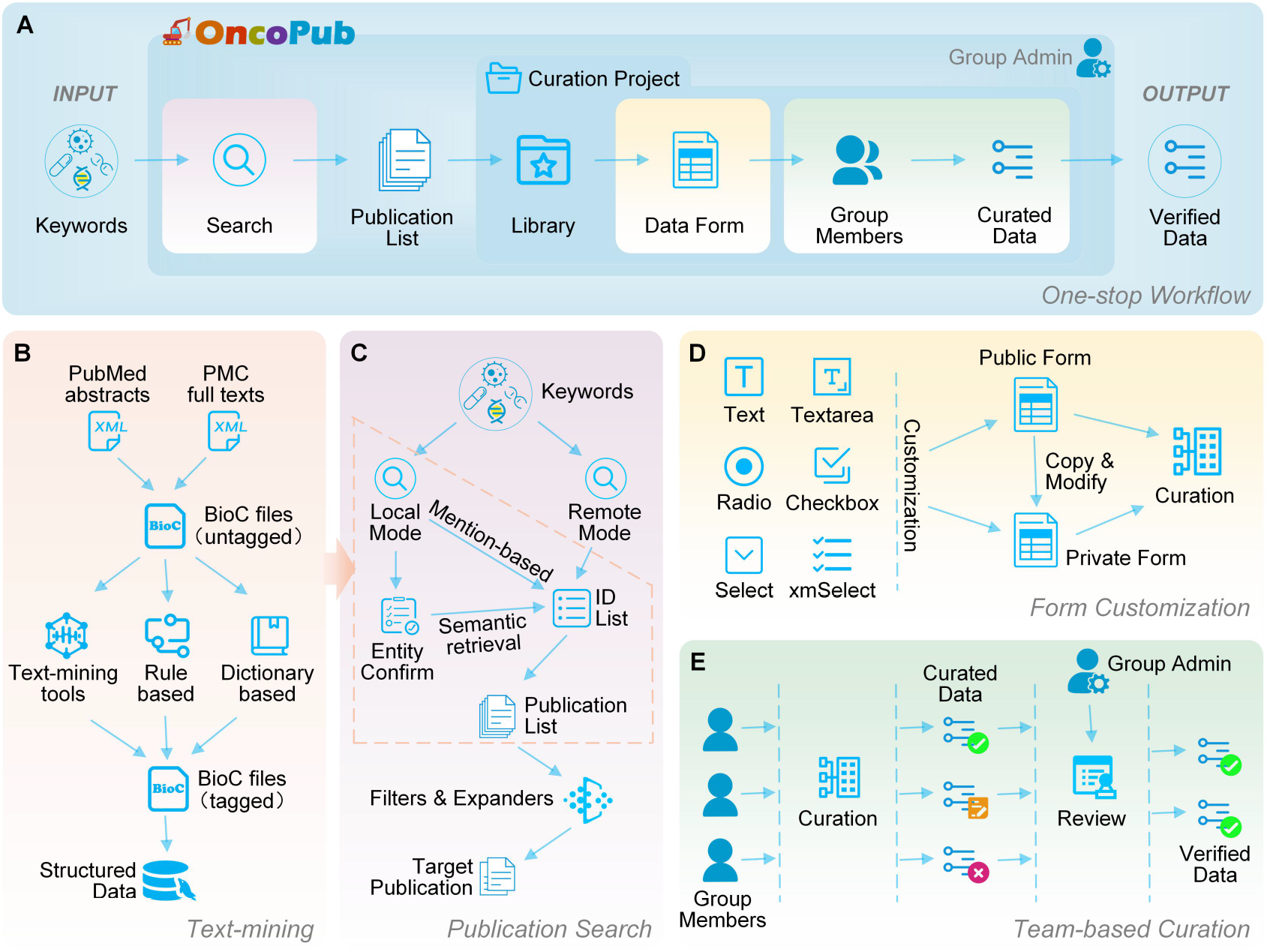
OncoPubMiner overview. The one-stop “keyword in, knowledge out” workflow of OncoPubMiner is presented in (A), the text mining methodology is shown in (B), and the design related to publication search, form customization, and manual curation in the workflow is demonstrated in (C), (D), and (E), respectively.

### Text Mining

#### Data Retrieval and Pre-Processing

The open-access PubMed abstracts and PMC full text articles were obtained from the National Center for Biotechnology Information FTP server. We first downloaded the baseline packages and then tracked the most recent updates of PubMed and PMC on a daily basis. All text in Extensible Markup Language format was transformed to BioC-JSON format, a community-driven biomedical text processing data format for improved interoperability, to simplify future data processing and exchange (36). Since sentences have a higher level of localization and information density than paragraphs, they are more likely to be relevant if they contain multiple bioentities (29). The Natural Language Toolkit (https://www.nltk.org/) is used to break down the transformed paragraphs into sentences.

#### Entity Recognition and Standardization

The system identified four primary categories of biological entities: disease/cancer, gene, alteration and chemical/drug. Here, we use DNorm (20), GNormPlus (21), tmVar (22), and tmChem (23) to mine disease, gene, alteration and chemical entities respectively. The software standardized the entities while mining, but this is not always the case. Moreover, the outcome of standardization of these software programs is to return only the identifier of the matched entry rather than the standard entry name. We continued to process these standardized results with scripts, obtaining standard entries while enhancing standardization. Medical Subject Headings (MeSH) is the standard used for cancer and drugs, while HUGO Gene Nomenclature Committee is for genes. We also map diseases to Disease Ontology (37) terms whenever possible to make them more interoperable with databases like CIViC (8). We adopted the standardized findings of tmVar without additional processing due to the vast number of variations and the lack of a single standard library akin to the MeSH entry library. Entity identification and term standardization provided three functions: providing the groundwork for the later creation of publishing retrieval services; pinpointing the position easier by highlighting relevant elements in retrieval results and publication reading; utilizing standardized entries to collect standardized data. Furthermore, the cancer entity tagging result categorized all publications as cancer-related or irrelevant, providing an additional filtering strategy for future literature searches.

One feature that separates OncoPubMiner from other text-mining systems is that it mines clinical significance, which defines how an alteration is connected to a certain clinical interpretation as described in the evidence statement, and evidence direction, which shows whether the evidence statement supports or refutes the clinical significance of an event. The tagging of these types of entities can assist users in swiftly locating crucial information to extract relational data, thereby considerably improving the extraction efficiency of publication data.

### Publication Search

OncoPubMiner provides two retrieval options for literature search: local retrieval based on entity annotation and remote retrieval based on PubMed E-utilities application interface (API). The local search mode was further divided into mention- and entity-based searches (see Fig. 1B for details). The mention-based search will return the articles whose text exactly or partly matches user’s input term. Entity-based search is a type of semantic search in which a keyword may match numerous standard phrases by fuzzy keyword matching. To increase the accuracy of the search results, this procedure is separated into two steps. The user’s keywords are utilized in the backend to search the standard entries, the matched entries are provided to the user, and the user selected entry is used to associate the article to retrieve the final article list.

OncoPubMiner takes several steps to perform a search operation. First, it queries the database to retrieve the publication’s identifier (PubMed ID). Second, the API is used to acquire the BioC-JSON data associated with the article from the OncoPubMiner server. Each publication, as previously stated, has two versions of BioC-JSON: one after the original data has been preprocessed and transformed, and another after the texts have been tagged. The two BioC-JSON versions are separated by a temporal gap. As a result, the retrieved articles may have data but no entity labeling information, and such articles may lack a tag sign after the title when compared to the labeled ones.

### Data Collection Form

#### Form Customization

OncoPubMiner allows users to establish online data models (Fig. 2). As a public platform, data form customization is essential for meeting the demands of different institutions for one-stop publication data collection. OncoPubMiner allows users to build and modify data collection forms. Users can customize the name, type, prompt text and position of each item in the form, and to specify whether the item is required. Users can customize the options (for radio/checkbox and select types) but use standard cancer types, genes, and drug libraries pre-integrated in the system as the option list (xmSelect type). The maximum length for text-type form items (text and text area) and the maximum quantity for multiple-selection form items (checkbox and xmSelect) can be specified. Furthermore, whether the field is mandatory, as well as default values or default options, can be specified. Users can use terms such as “[PMID]” and “[TITLE]” as default values in text type fields (text or text area) to allow automated collection of article ID and article title (see Supplementary File 2, Figs. 22– 23). The system will automatically fill in the required information from the current literature during the subsequent data collection. All of these parameters and constraints can help guarantee that curators collect data in line with the desired format and standards, resulting in standardized and structured knowledge data.

**Figure 2.**
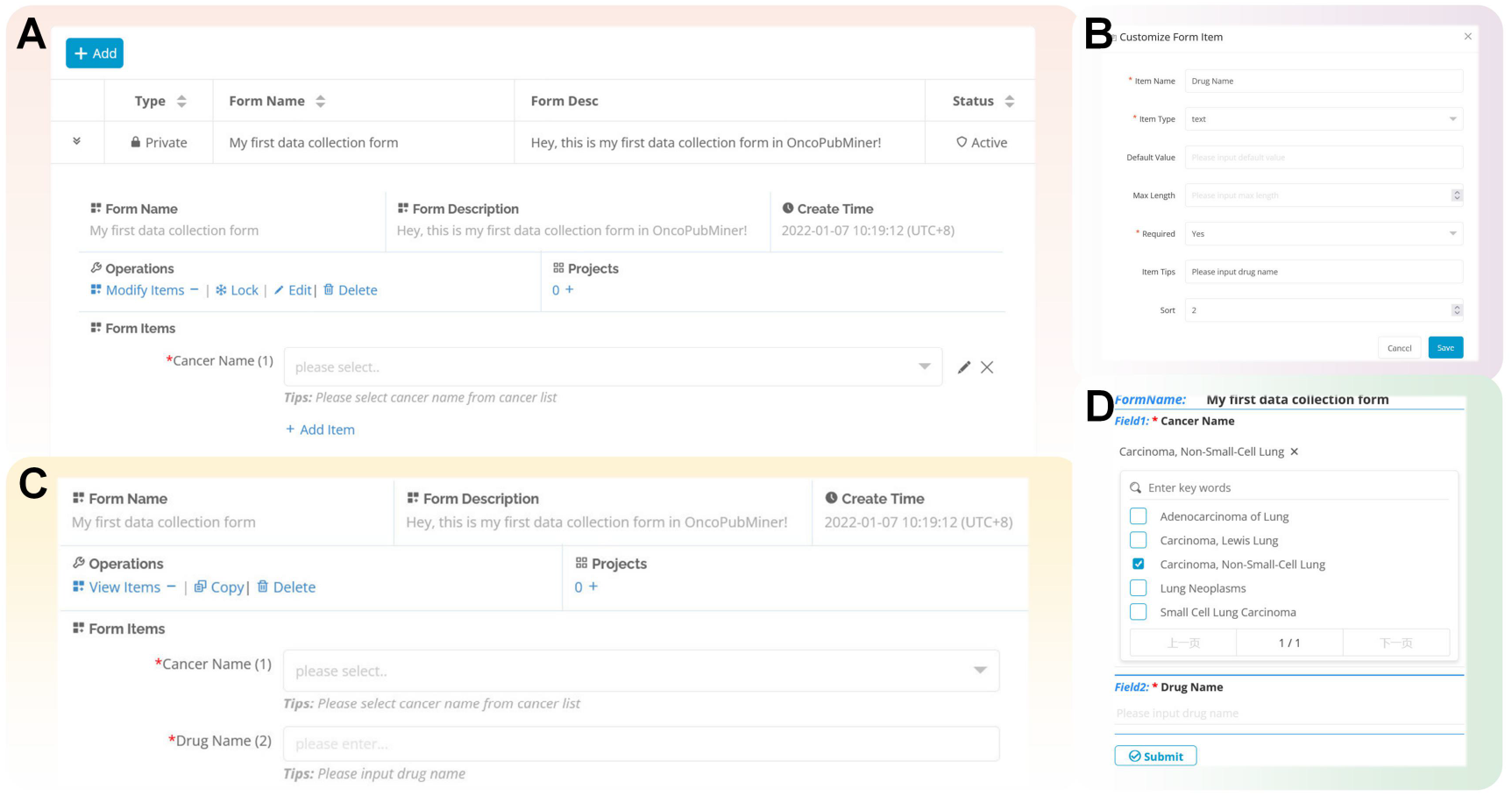
Data collection form customization. (A) Newly created data collection form. (B) Form item definition. (C) The status-locked data collection form. (D) The locked form is ready for data collection.

#### CIViC Data Forms and Feedback

CIViC is a community knowledgebase for expert crowdsourcing the clinical interpretation of variants in cancer. Since its initial public release in 2017, CIViC has gained widespread recognition and been used as a result of its entirely open features, free community collaboration platform, and high-quality data. It is expected to play a significant role in the tumor precision diagnostic and treatment community when combined with its peer-reviewed standard operating procedure (SOP) (38). To interact with this open platform, OncoPubMiner predefines a set of standardized data collection templates that adhere to the CIViC SOP, which can be used by all users to extract data from the literature. Because the acquired data adhere to the CIViC data standard, it is simple to connect with the CIViC database to enable quick data upload and exchange. These data forms may also be used as templates for users to copy, edit, and improve upon, allowing users to easily construct the data model they require.

CIViC provides data download features for all five categories of evidence as an opensource knowledge base platform. We obtained the nightly data files from CIViC (https://civicdb.org/releases), and after simple processing, the evidence data were incorporated into the OncoPubMiner knowledge base in the format of the above-mentioned pre-integrated forms. The data will be downloaded, parsed, and imported into the knowledge base on a monthly basis, and the new data will completely overwrite the old data. In the latest version of the system (2022-01-01), a total of 3,843 pieces of evidence have been incorporated, with 510, 643, 2,403, 67, 27, and 193 of these being prognostic, predisposing, predictive, oncogenic, functional, and diagnostic. These data may be placed on the associated literature page, and visitors can understand how each piece of CIViC evidence was gathered intuitively.

### System Implementation

OncoPubMiner is divided into the application system (APP) and APIs. The SpringBoot (version 2.3.1) and Mybatis-plus (version 3.3.2) frameworks were used to create the OncoPubMiner APP. LayUI (version 2.5.6), EasyWeb (version 3.1.8), and jQuery (version 3.2.1) were used to build the front end. OncoPubMiner APIs provide users with access to all publication search features from a programming environment in the BioC-JSON based format. The APIs were written in Python and built with the Flask-RESTful framework. Both the APP and APIs were supported by MySQL as the database management system. Additionally, much effort was devoted to improve the systems to be compatible with browsers in mobile devices.

## USAGE

OncoPubMiner can be accessed through a user-friendly web interface. The platform can be used as a publication search system with a variety of retrieval modes and filtering methods, or as a one-stop KI-KO platform for extracting structured formats of knowledge sets triggered by certain keywords. OncoPubMiner has several advantage over the existing similar systems (Supplementary File 1: Table S1).

### Publication Search and Library Management

#### Publication Search

To search for publications, the user must enter the keywords and specify the necessary parameters (Fig. 3A). The search results page is divided into left and right columns. The search or display parameters are located on the top left, and all entities recognized from the literature are located below. The publications are shown on the right with title, authors, journal information, and abstract.

**Figure 3.**
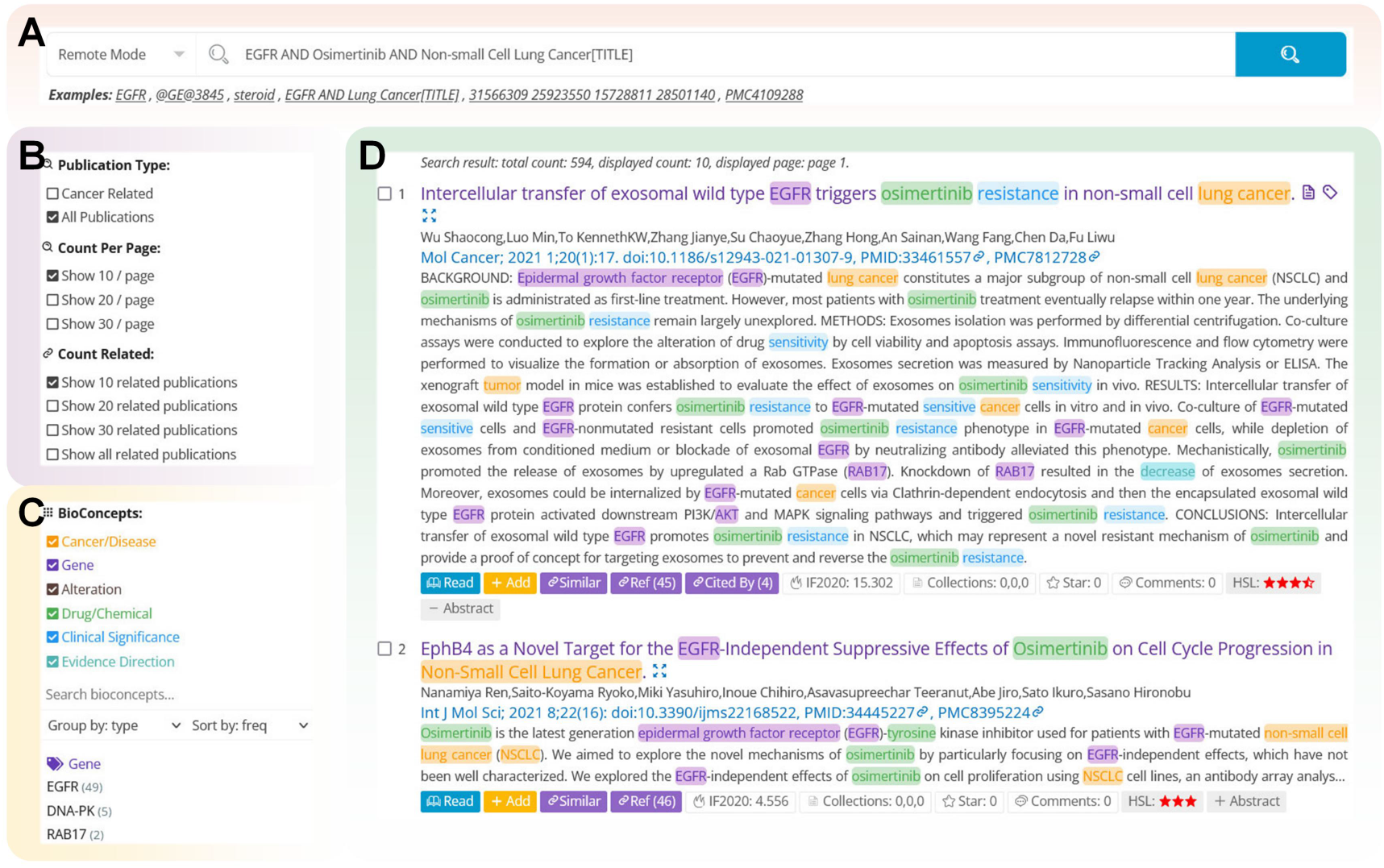
Publication search page. (A) Publication search panel. (B) Parameters setting panel. (C) Bioconcepts identified from displayed articles. (D) Search results.

Through a keyword search, whether mention- or entity-based, many articles are obtained. Expert curation is needed to select best articles. To this end, OncoPubMiner examines the value of each article and determines whether it is curatable from several viewpoints. To aid curators in determining the article value, we show the journal’s impact factors, the status of users’ rating and commenting on articles, and the status of articles being included in the collection. Notably, we established the highest-sentence level scoring system to score the amount of distinct entity categories occurring in the sentence of the article (see Table S3 in Supplementary Material 1 for details) to help curators estimate an article’s value, hence assisting curators by streamlining the literature triage process. Having similar findings from independent studies is common in research and is important to assess reproducibility (14), OncoPubMiner provides citations and referenced papers connected to the retrieved articles, as well as similar publications, which can assist in rapidly expanding the scope of publications.

#### Library Management

OncoPubMiner can help manage the retrieved publication after search. Users may set up a publication collection on the library management page. When creating a publication collection, the user need to input the name and description (optional), which may be used to differentiate among collections. Users can maintain the keyword lists concurrently if the collection exclusively focuses on publications linked to certain keywords or phrases, and these keywords will be available for local or remote retrieval of articles to rapidly explore current collection relevant ones. The user can only add publications to the collection once it has been created and locked.

Users may search for articles by clicking on the link of a specified set of keywords on the library page. They can also manually enter terms into the literature search page to conduct a search. Users can add a single publication to the search list by clicking the ‘Add’ button behind each one in the list. When the addition is finished, the corresponding literature will be available on the library page. It should be noted that the same article might be included in multiple collections to achieve various curation aims.

### Biocuration Projects

#### Team Account Management

OncoPubMiner is a free system, with features such as publication retrieval, viewing, and data collection available to everyone. However, because data collection for knowledge bases, particularly those related to precision diagnosis and treatment of malignancies, has such strict quality standards, data quality must be assured through collaborative team evaluation. As a result, OncoPubMiner has developed a collaborative review mode. In this mode, when a user first creates an account, a group is also created with this user as the group administrator, which can create accounts for team members. Users do not need enter any personally identifiable information (e.g., name, email, institution). The account is only required to assist the system in managing various data as a team to ensure data security.

#### Biocuration Projects

The biocuration process should be project-centric, focusing on the administration of a certain kind of mining objective such as biocuration-related curators, data structures, to-be-mined publication list, and eventually mined data. Users may build projects on their own and with team members. To prevent future data discrepancies caused by modifications, the data collection form and publication collection must be locked before being attached to a curation project. Except for the publication related to the collection, which may be continually updated, all aspects in the locked project cannot be modified.

#### Literature Curation

OncoPubMiner provides online reading, publication curation, and data review, which as a true literature curation platform that generates high-quality knowledge data. After the project has been created and locked, team members are able to visit it under their account. The user must expand the project associated publication table and click on the ‘Read’ link after each publication to enter the curation page for publication reading and data extraction operations (Fig. 4). The page is divided into three parts. In the middle is the paper viewer, which displays the annotated publication with highlighted entities. The full-text article of PMC will be displayed if available; otherwise, the abstract will be displayed. The articles are shown in the form of sentences, and the level of the sentence is displayed in the shape of red stars in front of each sentence. The more stars, the more essential the statement, making it easier for the curator to find the critical location. The left side of the page presents a list of entities identified from this document, which can be grouped and sorted in different ways. The right side of the page reveals the form linked with the present project, which is more essential. All form items are presented one-by-one in the order specified. Users may extract data from the page viewer, edit each field, and submit it after review and approval. After all members complete the data submission and mark the article as complete, the group administrator can start the data review, ensuring that the data is high quality.

**Figure 4.**
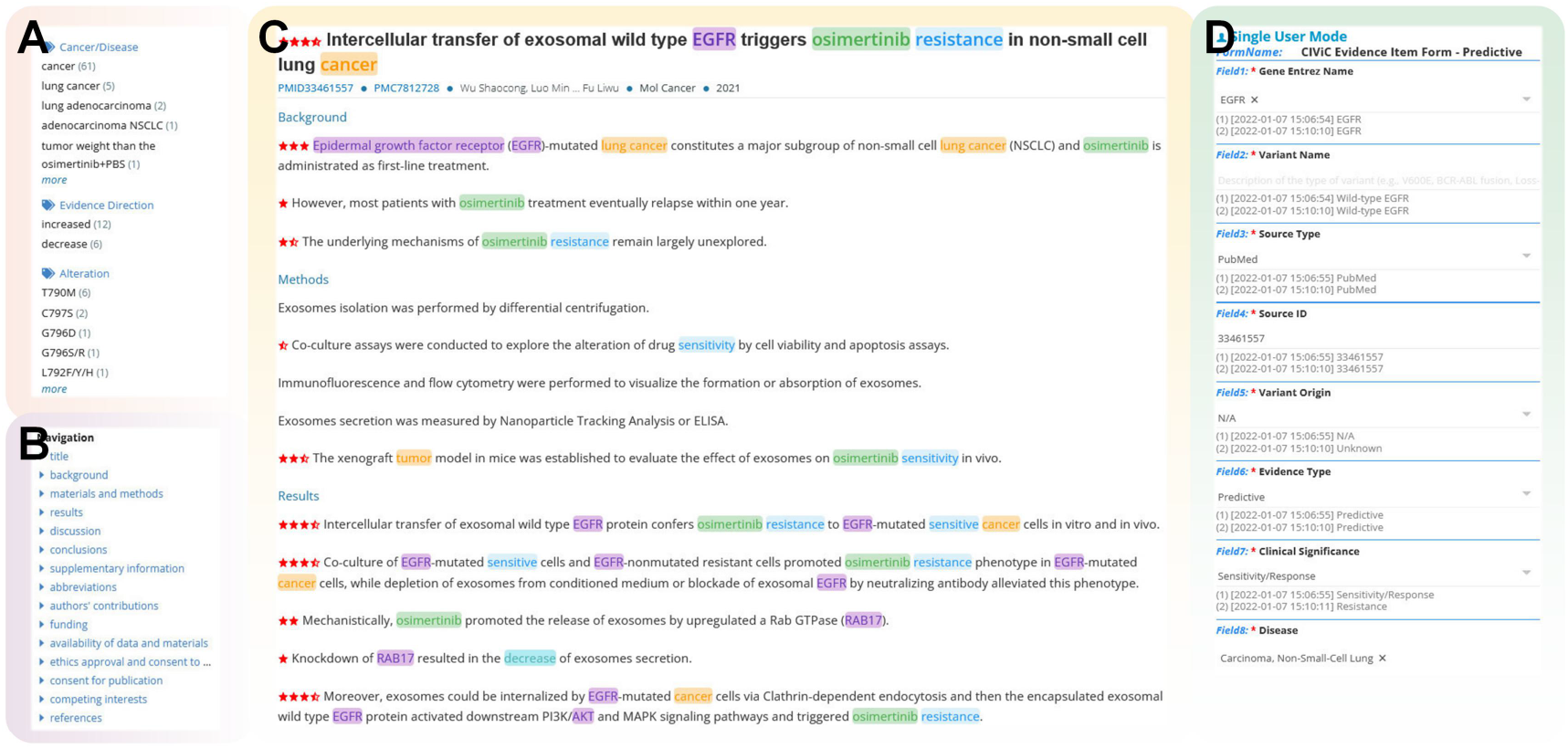
Literature curation page. (A) Bioconcepts identified from the article. (B) Navigation for each section of the article, which appears only when the full PMC text is available. (C) The text of the article with the annotated entities highlighted. (D) Data collection panel.

There is a need for high-quality data for constructing knowledge bases, but there are also users who need to rapidly read the literature and extract data. OncoPubMiner has a single-user mode for literature curation to satisfy such short data collection demands. The user hits the “Read” button behind the item on the publishing list page of the search results, and then selects the data collection form to be utilized in the pop-up box to access the details page, as shown in Figure 4. When data collection is complete, the data may be visible and downloaded right away. This mode also allows numerous sets of data for the same article to be submitted and downloaded and allows both logged-in and non-logged-in users to read literatures and submit data. Of course, the user is also able to just read literature without selecting data.

### User Guide

We provide a user guide (see Supplementary File 2) that covers all the key features of the system, including publication search, library management, account and team management, data collection form customization, curation project management, among others. The examples used in this user guide are available through the demonstration accounts, which include a group administrator and three team members. Any user may log in and access this information by clicking on any account link and find step-by-step tutorials.

## CONCLUSIONS

In this paper, we present OncoPubMiner, a free platform for mining oncology publications. The study combines text mining, data structure customization, article search with online reading, project-centered and team-based data collection to realize a one-stop KI-KO data collection system. All open-access PubMed abstracts and PMC full-text articles are downloaded and updated on a daily basis, and all of the raw publication data retrieved is transformed to BioC-JSON format. The retrieved literature is managed by the library and may be linked to the curation project for team-based literature data extraction and evaluation. Since the system allows customizing of data collection forms online, any group might mine the literature on cancer precision medicine and generate structured knowledge data to build knowledge bases or develop bioinformatics applications. OncoPubMiner uses NLP technology to perform entity recognition on the combined PubMed and PMC contents. In addition, although focused on precision oncology, the generalizability of the data structure customization feature and the system’s team collaborative review mode can assist with gathering various types of information such as drug-drug, drug-gene interactions, and gene- cancer relationships. If the entity recognition step allows for the tagging of many more categories of biological entities, OncoPubMiner’s use will undoubtedly become considerably broader.

In the future, we will deliver better entity recognition algorithms, software, and terminology databases to recognize new types of entities and improve the tagging results of existing entity categories. In addition, we plan to provide relationship extraction capability, which will allow for the extraction of entity connection information. Combined with the current work process, these modifications will significantly improve efficiency with quality assurance. Under OncoPubMiner’s support, we may generate more consistently updating expert-quality knowledge bases.

## AVAILABILITY

OncoPubMiner is free and open to all users. OncoPubMiner can be accessed at https://oncopubminer.chosenmedinfo.com.

## ACCESSION NUMBERS

None

## SUPPLEMENTARY DATA

Supplementary Data are available at NAR online.

## ACKNOWLEDGEMENT

We thank LetPub (www.letpub.com) for their linguistic assistance during the preparation of this manuscript.

