## Supplemental File 1 for "OncoPubMiner: A platform for oncology publication mining"

**Supplementary Table S1: Comparison of OncoPubMiner to other systems.**

| Features / Systems | OncoPubMiner | BEST | GeneView | LitSense | LitVar |
|---|---|---|---|---|---|
| **Text-mining** | | | | | |
| Contents | PubMed + PMC | PubMed | PubMed + PMC | PubMed + PMC | PubMed + PMC |
| Entity recognition | Public accessed tools + Self build dictionary based lookup | Dictionary-based approach | Public accessed tools + Dictionary-based approach | PubTator | Public accessed tools |
| Entity types | Cancer/Disease, Gene, Alterations, Drug/Chemical, Clinical significance, Evidence direction | Gene/Protein, Target, Transcription Factor, miRNA, Chemical Compound, Drug, Toxin, Disease, Pathway | Genes, SNPs, Species, Chemicals, Histone modifications, Cell types, Diseases, Drugs, Enzymes, Tissues | Gene, Mutation, Disease, Chemical, Species, Cell Line | Genes, Chemicals, Diseases, Species |
| **Publication Search** | | | | | |
| Update frequency | Daily (twice a day) | Daily | Quarterly | Regularly | Monthly |
| Search Mode(s) | Local entity-based matching, Local mention-based matching, Remote E-utilities-based [illegible] | Keyword Query | Entity-specific search, Advanced keyword-based search | Sentence search | Three-step normalization process-based search |
| Entity confirm | √ | √ | - | - | √ |
| Importance Label | GitHub-like star, Comments, Journal impact factor, Highest sentence-level | - | - | - | - |
| Cross-link | Cited by, References, Similar | - | - | - | - |
| Entities Highlight | √ | √ | √ | √ | √ |
| Library management | Collections, Publications | - | - | - | - |
| RESTful API | √ | - | - | - | √ |
| **Manual Curation** | | | | | |
| Curation Mode(s) | Collaborative Mode + Single User Mode | - | - | - | - |
| Form customization | Supports customization of multiple types of | - | - | - | - |
| Protocol-based Forms | Currently supports CIViC data form protocol | - | - | - | - |
| Project management | √ | - | - | - | - |
| Sentence-level display | √ | - | - | - | - |
| Importance label | 10 grade star labels | - | - | - | - |
| Multiple review | √ | - | - | - | - |
| Data echo | CIViC data echo, Collected knowledge data echo | - | - | - | - |
| Data downloads | Form related structured data | - | Annotated results | - | Search results |
| Knowledge base oriented | √ | - | - | - | - |

(to be continued)

**Supplementary Table S1: Comparison of OncoPubMiner to other systems.**

(continue)

| Features \ Systems | PubTator Central | PubTerm | SciLite | TeamTat | Textpresso Central | Thalia |
|---|---|---|---|---|---|---|
| **Text-mining** | | | | | | |
| Contents | PubMed + PMC | PubMed | Europe PMC | PubMed + PMC, User-documents | WormBase C. elegans bibliography, | PubMed |
| Entity recognition | Public accessed tools | PubTator | Dictionary-based approach + machine-learning based filter | Public accessed tools | Named Entity Recognizers | Public accessed tools |
| Entity types | Gene, Variant, Disease, Chemical, Species, Cell Line | Genes, Diseases, Chemicals, Species, Sequence variants | Gene/Protein, Organisms, Diseases, GO terms, Chemicals, Gene function, Phosphorylation event | Chemical, Disease, Gene, Species, Variation | Gene Ontology, Sequence Ontology, Chemical, Phenotype and Trait Ontology, Protein Ontology, Species | Chemicals, Diseases, Drugs, Genes, Metabolites, Proteins, Species, Anatomic |
| **Publication Search** | | | | | | |
| Update frequency | Daily | Monthly | Daily | Uploaded by user | Monthly | Daily |
| Search Mode(s) | Keyword search, Semantic search | PubMed Query | Europe PMC | - | Lucene-based search | Entity-based matching |
| Entity confirm | - | - | - | - | - | √ |
| Importance Label | - | - | - | - | - | - |
| Cross-link | - | - | - | - | - | - |
| Entities Highlight | √ | √ | √ | √ | - | √ |
| Library management | Collections | - | - | - | - | - |
| RESTful API | √ | - | - | - | - | √ |
| **Manual Curation** | | | | | | |
| Curation Mode(s) | - | - | - | Collaborative Mode + Single User Mode | Single User Mode | - |
| Form customization | - | - | - | - | √ | - |
| Protocol-based Forms | - | - | - | - | - | - |
| Project management | - | - | - | √ | - | - |
| Sentence-level display | - | √ | - | - | - | - |
| Importance label | - | - | - | - | - | - |
| Multiple review | - | - | - | √ | - | - |
| Data echo | - | Annotated results | - | Annotated results | Annotated results | - |
| Data downloads | - | Annotated results | - | Annotated results | Annotated results | - |
| Knowledge base oriented | - | - | - | - | √ | - |

**Supplementary Table S2: The dictionary for recognizing clinical significance and evidence direction concepts.**

| Concept type | Entity category | No. | Entity | Aliases |
|---|---|---|---|---|
| Clinical significance | Predictive | C1-01 | Sensitivity | Sensitive; Sensitiveness; Sensibility |
| | | C1-02 | Insensitivity | Insensitive |
| | | C1-03 | Response | Responsive; Efficacy |
| | | C1-04 | Resistance | Resistant |
| | | C1-05 | Adverse event | ADR*; Adverse reaction; Adverse Drug Reaction; Adverse Drug Event |
| | | C1-06 | Toxicity | Cytotoxicity |
| | | C1-07 | Predict | Predictive |
| | Diagnostic | C2-01 | Positive | / |
| | | C2-02 | Negative | / |
| | | C2-03 | Diagnosis | Diagnostic |
| | Prognostic | C3-01 | Outcome | / |
| | | C3-02 | Prognosis | Prognostic |
| | | C3-03 | Survival | CR*; PR*; SD*; PD*; DFS*; OS*; PFS* |
| | | C3-04 | ORR* | / |
| | | C3-05 | DCR* | / |
| | Functional | C4-01 | Function | Functional |
| | Predisposing | C5-01 | Predisposing | Predispose |
| | | C5-02 | Risk | / |
| | | C5-03 | Susceptibility | Susceptible; Susceptibilities |
| | General | C6-01 | Marker | / |
| Evidence direction | Up | D1-01 | Gain | / |
| | | D1-02 | Increased | Increase; Increasing |
| | | D1-03 | Better | / |
| | | D1-04 | Support | Supports |
| | | D1-05 | Activate | Active |
| | Down | D2-01 | Poor | / |
| | | D2-02 | Inactivate | Inactive |
| | | D2-03 | Loss | / |
| | | D2-04 | Does not support | / |
| | | D2-05 | Decreased | Decrease; Decreasing |
| | / | D3-01 | Unaltered | Unchanged |
| | | D3-02 | Unknown | N/A; NA*; Not Available |

* Case Sensitive (Otherwise, case insensitive)

**Supplementary Table S3: The criteria for assigning a grade to each sentence in the publication.**

| No. | Sentence-level | Criteria Description |
|---|---|---|
| 1 | ☆ | Clinical significance |
| 2 | ★ | Cancer (+ Evidence direction) / Gene (+ Evidence direction) / Alteration (+ Evidence direction) / Drug (+ Evidence direction) / Clinical significance + Evidence direction |
| 3 | ★☆ | Cancer + Clinical significance / Gene + Clinical significance / Alteration + Clinical significance / Drug + Clinical significance |
| 4 | ★★ | Cancer + Gene (+ Evidence direction) / Cancer + Alteration (+ Evidence direction) / Cancer + Drug (+ Evidence direction) / Cancer + Clinical significance + Evidence direction / Gene + Alteration (+ Evidence direction) / Gene + Drug (+ Evidence direction) / Gene + Clinical significance + Evidence direction / Alteration + Drug (+ Evidence direction) / Alteration + Clinical significance + Evidence direction / Drug + Clinical significance + Evidence direction |
| 5 | ★★☆ | Cancer + Gene + Clinical significance / Cancer + Alteration + Clinical significance / Cancer + Drug + Clinical significance / Gene + Alteration + Clinical significance / Gene + Drug + Clinical significance / Alteration + Drug + Clinical significance |
| 6 | ★★★ | Cancer + Gene + Alteration (+ Evidence direction) / Cancer + Gene + Drug (+ Evidence direction) / Cancer + Gene + Clinical significance + Evidence direction / Cancer + Alteration + Drug (+ Evidence direction) / Cancer + Alteration + Clinical significance + Evidence direction / Cancer + Drug + Clinical significance + Evidence direction / Gene + Alteration + Drug (+ Evidence direction) / Gene + Alteration + Clinical significance + Evidence direction / Alteration + Drug + Clinical significance + Evidence direction |
| 7 | ★★★☆ | Cancer + Gene + Alteration + Clinical significance / Cancer + Gene + Drug + Clinical significance / Cancer + Alteration + Drug + Clinical significance / Gene + Alteration + Drug + Clinical significance |
| 8 | ★★★★ | Cancer + Gene + Alteration + Drug (+ Evidence direction) / Cancer + Gene + Alteration + Clinical significance + Evidence direction / Cancer + Alteration + Drug + Clinical significance + Evidence direction / Cancer + Gene + Drug + Clinical significance + Evidence direction / Gene + Alteration + Drug + Clinical significance + Evidence direction |
| 9 | ★★★★☆ | Cancer + Gene + Alteration + Drug + Clinical significance |
| 10 | ★★★★★ | Cancer + Gene + Alteration + Drug + Clinical significance + Evidence direction |

**Supplementary Table S4: Elements for data collecting form item customisation.**

| No. | Element Name | Element Type | Value Type | Description | Required |
|---|---|---|---|---|---|
| 1 | Item Name | Input | String | The name for data collection form item, it will also serve as the field name of the final dataset. | Required. |
| 2 | Item type | Select | String | Including input, textarea, radio, checkbox, select and xm-select*. | Required. |
| 3 | Option Type | Select | String | Including customize, cancer list, gene list, alteration list and drug list. | Required for select and xm-select |
| 4 | Option List | Input list | String | Define the option list for item type radio and checkbox, and select or xm-select in customize option type. | Required for option type customize, or item type radio and checkbox |
| 5 | Default Value | Input | String | Define the default value for item type input or textarea. | Not required. |
| 6 | Default Value | Xm-select* | String | Define the default value for item type radio and checkbox, and select or xm-select in customize option type. | Not required. |
| 7 | Max Length | Input | Integer | Define the max length of the input text for item type input or textarea, default 0, means no length restriction. | Not required. |
| 8 | Max Count | Input | Integer | Define the max count that can be selected for item type checkbox and xm-select, default 0, means no length restriction. | Not required. |
| 9 | Required | Select | Boolean | Whether the item must be filled or selected when collection data, default not required. | Required. |
| 10 | Item Tips | Input | String | The tips to inform the users what this item means to, and/or something need to be paid attention | Not required. |
| 11 | Sort | Input | Integer | Define a numeric value to help sort the items in the sample data collection form, default 100. As for items with the same sort number, the create time of each item will be served as the sort standard. | Not required. |

* xm-select (https://gitee.com/maplemei/xm-select) is an open source multiple selection drop-down menu based on the open source front-end frame LayUI (https://github.com/sentsin/layui, https://gitee.com/sentsin/layui).
