## Supplemental File 2 for "OncoPubMiner: A platform for oncology publication mining"

**User Guide**

### 1. Library Management

The OncoPubMiner library is comprised of multiple collections that centrally gather articles on similar topics, and the library administration page is only available to group administrators (GAs), namely the accounts generated through the system registration function. GAs may create and manage collections on the 'Library' page, as well as search for and manage articles inside collections.

#### 1.1 Collection-Oriented Tab

The library administration website is divided into two tab pages: Collection-Oriented and Publication-Oriented. GAs may manage all collections on the Collection-Oriented tab page.

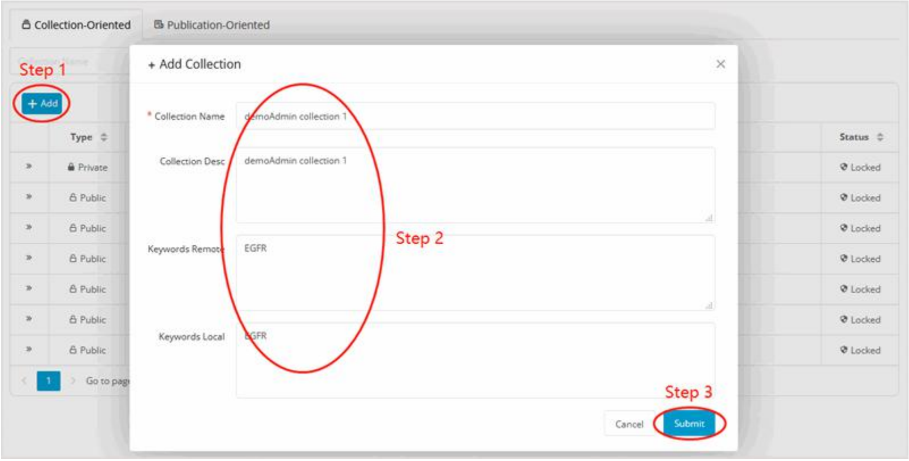

Figure 1. Create a new collection.

Click the ‘Add’ button, enter collection-related information in the pop-up box, such as the name, description, and a list of keywords for remote and local retrieval articles, and then click the ‘Submit’ button to finish the collection's creation (Figure 1).

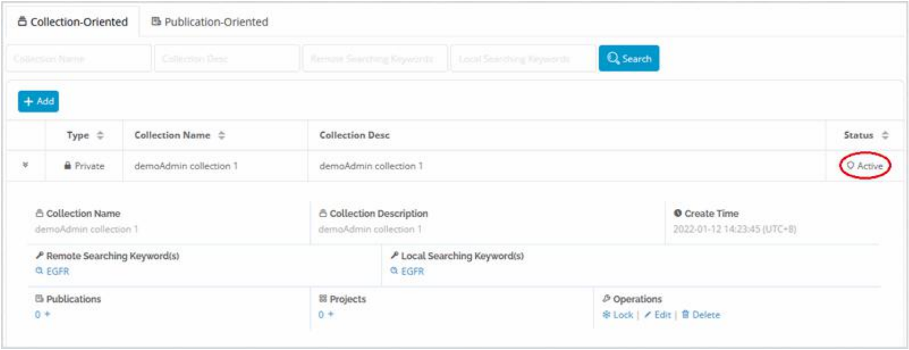

Figure 2. The newly created collection.

As illustrated in Figure 2, newly created collection is active by default, and all information is editable. Collection in this state is ineligible for article management, but it may be used for literature inclusion and project creation wherever the GA hits the lock link and successfully locks the collection. Because there are no associated projects, this collection can be deleted at any time.

When users click the keyword for a remote or local search, the system will open the literature search page and utilize the term to do the related literature search. If the current collection is locked, 1) the obtained publications may be added to the collection by GA, and 2) the current collection can be associated while the curation project is being created or modified.

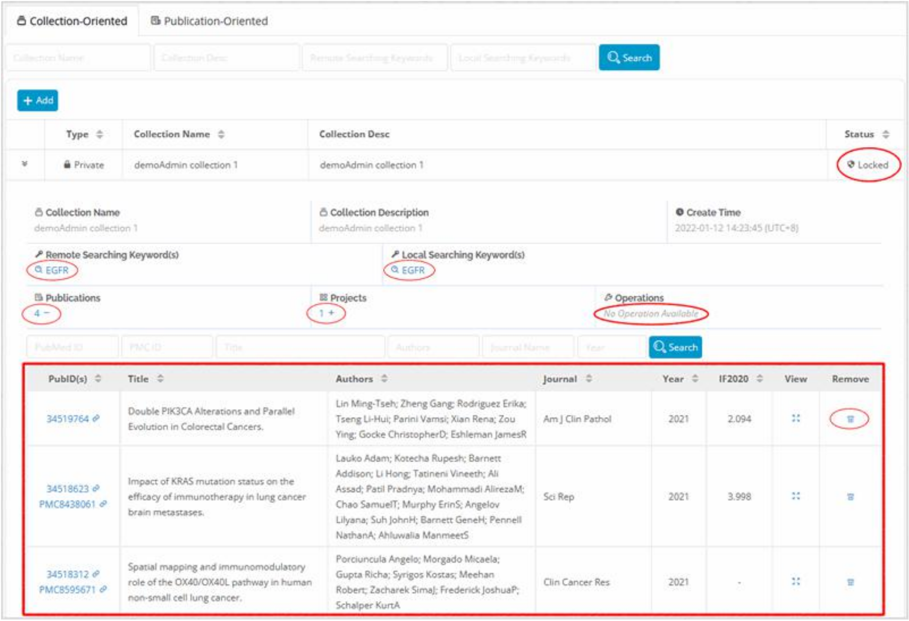

Figure 3. A locked collection that contains articles and is associated with the curation project.

Figure 3 depicts the interface of the locked collection after it has been associated with the curation project and the articles have been included. By selecting a number under text ‘Publications’ or ‘Projects’, you will be sent to a list of all articles or projects related with that collection. The article list displays the article's basic information as well as a link to the article details page. Furthermore, each item in this list has a ‘Remove’ link that may be clicked to remove the relevant article from the list. The project list will provide the basic information for all curation projects associated with the collection. If the project has gathered data, you may also see how much data has been gathered here, and then click the link to see or download the data (as shown in Figure 4). Additionally, because the collection is tied to the curation project, it can no longer be immediately destroyed.

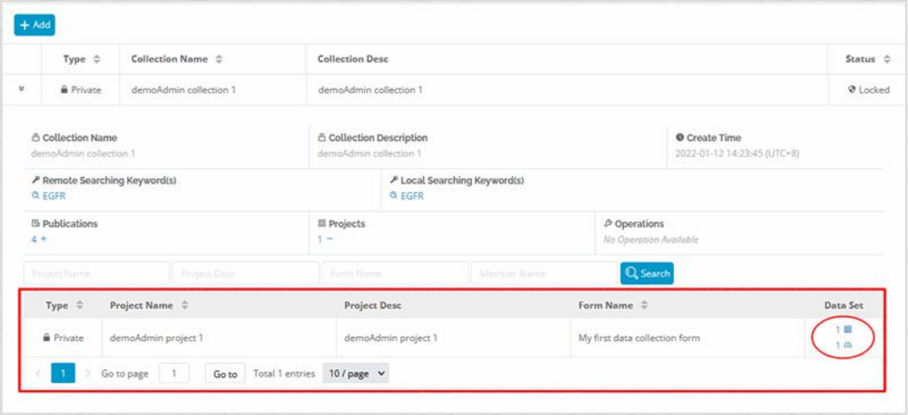

Figure 4. A locked collection is presented, along with the accompanying curation project table.

### 1.2 Publication-Oriented Tab

The Publication-Oriented tab page displays a list of all non-redundant articles in the collections of the current user group (see Figure 5). Users may simply search through all of the included literature by PubMed ID, PMC ID, title, authors, or journal.

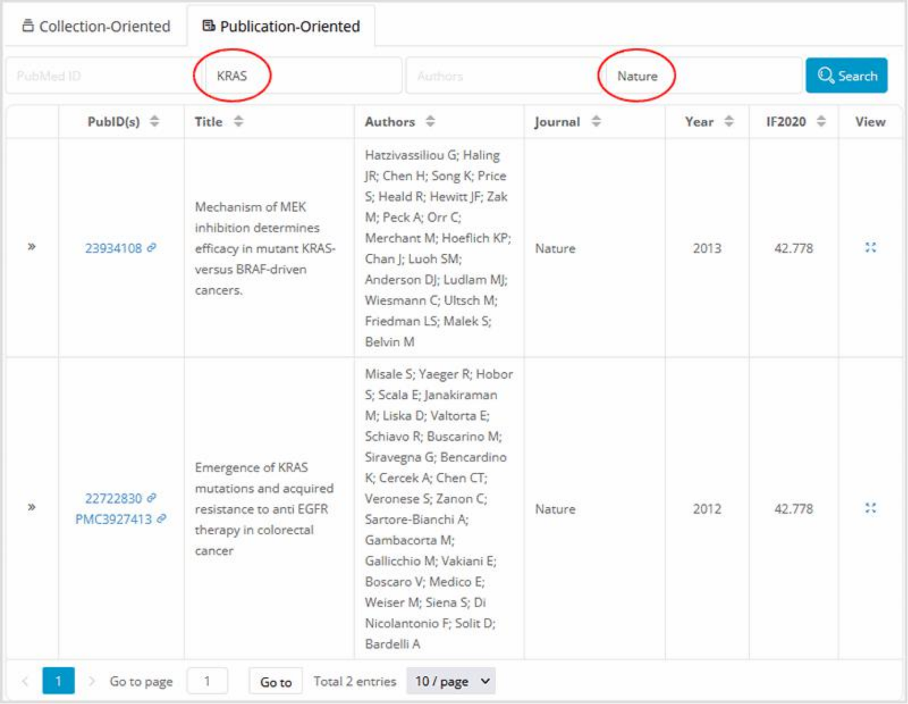

Figure 5. Publication-Oriented tab page.

### 2. Account Management

OncoPubMiner is a fully free tool for analyzing tumor-related literature. It may be used with or without a login information. However, in order to protect the security of user data and fully utilize the system's collaborative working mode, we provide account management capabilities such as account registration and team member managing.

#### 2.1 Account Registration

Users can create their own account by filling out the information on the user registration page (https://oncopubminer.chosenmedinfo.com/register). As previously stated, the objective of creating an account is to better explore the system's functionalities and protect the user's data security. Account registration only requires the user to enter the most basic account and password information, and does not require the user to enter any information that may reveal personal privacy, such as institute or email address (as shown in Figure 6). There are no hard requirements for account or password settings, and users can basically set them at their leisure.

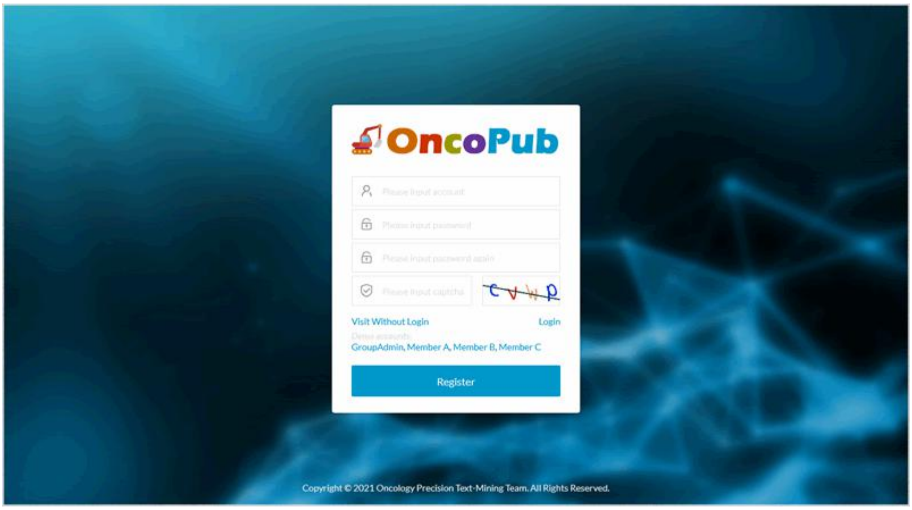

Figure 6. User registration page.

Users who have completed account registration can use their account and password to log in to the system via the login page (as shown in Figure 7). If the user does not have a registered account or is hesitant to establish one, he can log in with the relevant account by clicking the link of the demo account on the registration or login page. The system provides four demo accounts, which are the group administrator (group admin) and the team members (member A, B, and C) within the same user group.

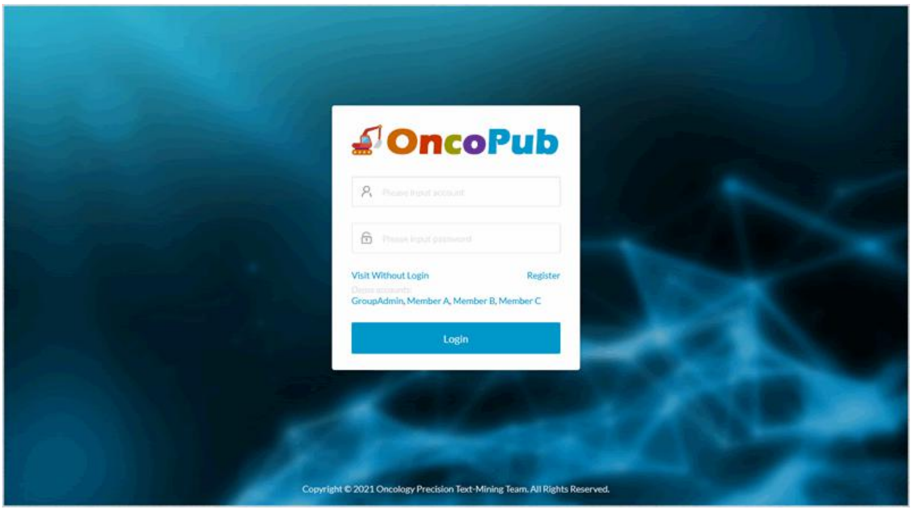

Figure 7. Login page.

**2.2 Team member management**

The account created through the user registration function is the administrator under the same name user group by default. After logging in to the system, he can enter the team member management page to create a member account for his team. As shown in Figure 8, after entering the team member account management page, the group administrator should click the ‘New Member’ icon, input the member account and password in the pop-up box, and then click the ‘Create’ button to finish the member account creation. Because freshly minted member have not yet joined any curation projects, its account is subject to deletion at any time (see Figure 9). However, it cannot be erased if it is related with the project. GAs can change member account passwords at any time, whether or not they are affiliated with a project.

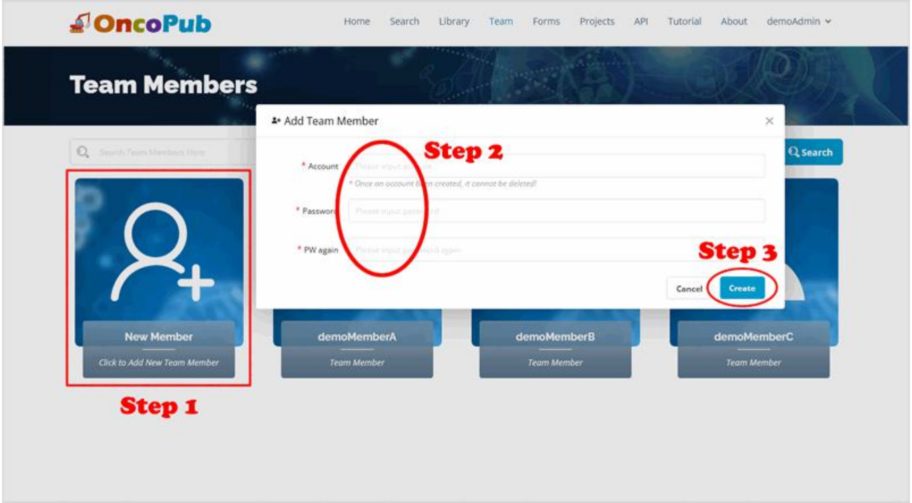

Figure 8. Account management page for team members.

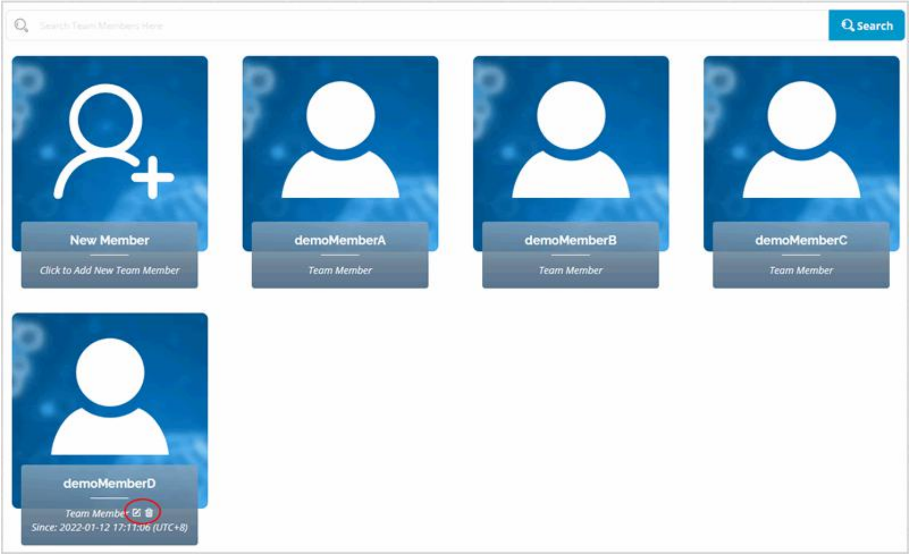

Figure 9. Newly established member accounts can be deleted at any moment.

It should be noted that demoMember A, B, and C in Figure 9 do not display the corresponding edit and delete icons not only because they are currently participating in a curation project, but also because the three member accounts are the demo accounts pre-created by the system, and in order to prevent malicious users from deleting these accounts or changing their login passwords after logging in to the system with demo GA account, the system does some special processing on these accounts and does not allow them to be edited or deleted.

### 3. Form Customization

Data collection forms may be established in the same way as article collections can, as seen in Figure 10.

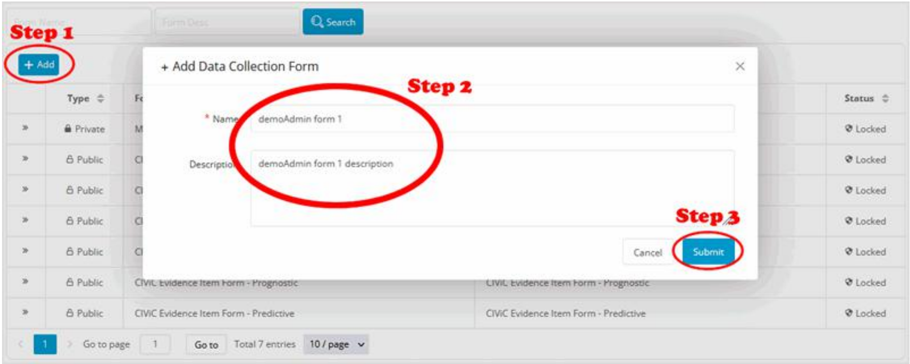

Figure 10. Create a new data collection form.

Figure 11 depicts the newly created data collecting form. It is active and may be modified, deleted, or locked at any moment. Because no form items have been created, the ‘Form Items’ section is empty. When you click the ‘Add Item’ link in this area, a pop-up window similar to the one shown in Figures 13 to 19 will appear, allowing you to customize the form item.

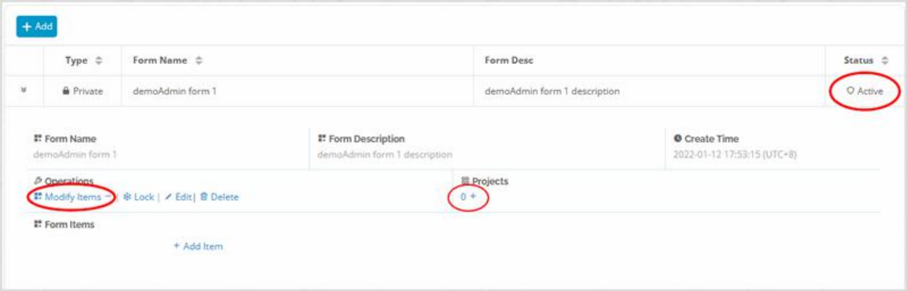

Figure 11. Newly created data collecting form.

OncoPubMiner supports the following form item types: text, textarea, radio, checkbox, select, and xmselect (see Figure 12). The first two are of the text category, and content must be provided manually, whereas the latter are of the option type, and alternative options can be specified in advance. Distinct form item types have different parameters that may be configured, which will be discussed in more detail below.

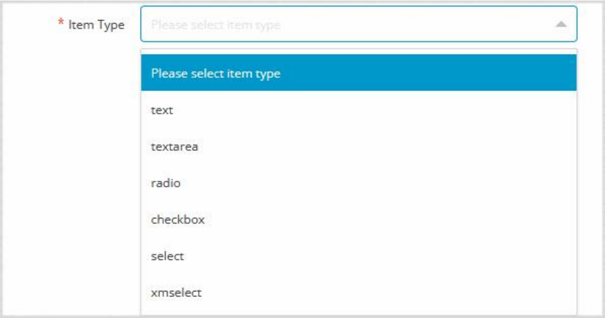

Figure 12. Options for form item type.

Item type 1: text

As shown in Figures 13–19, parameters such as item name, item tips, whether it is required, and the sort value used to order the form items are the same for any type of form item. The item name and tips are used to inform the user about the type of data collected by the field. Furthermore, the former will be utilized as the field name of the final data. Whether it is required determines whether a non-null judgment is made on the current field on the data collecting page. Required items do not accept missing value, which is critical for ensuring the integrity of essential field data.

In addition to the fields listed above, the text type item has the ability to set the default value as well as the maximum length of the field (see Figure 13). These two fields are, of course, optional. The default value is blank, indicating that no default value has been defined. The maximum length is blank, indicating that the field length is unrestricted.

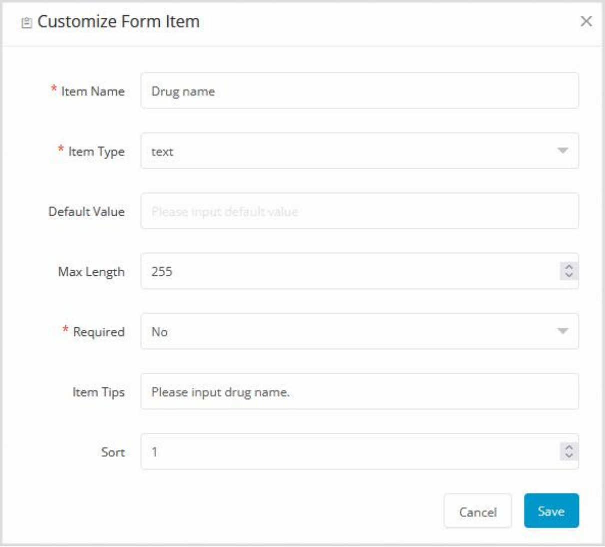

Figure 13. Customization of a text type form item.

Item type 2: textarea

The parameter settings for the textarea's item type (Figure 14) and the text kind are identical.

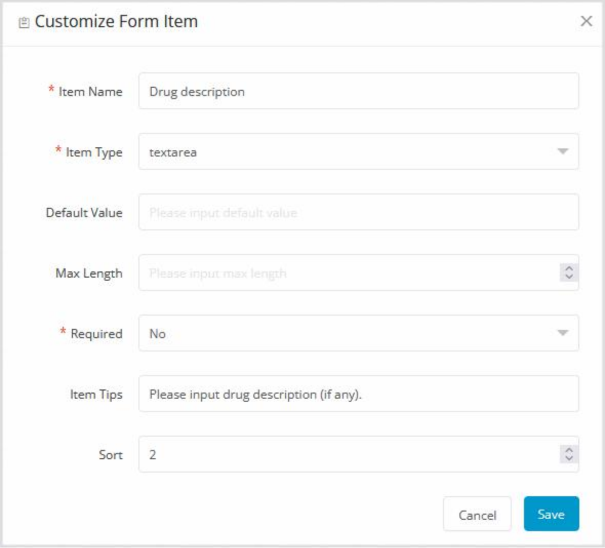

Figure 14. Customization of a textarea type form item.

Item type 3: radio

GAs can preset option values for the item type of radio type and select one of these preset option values as the default value (Figure 15). The default value, like the text type, is optional.

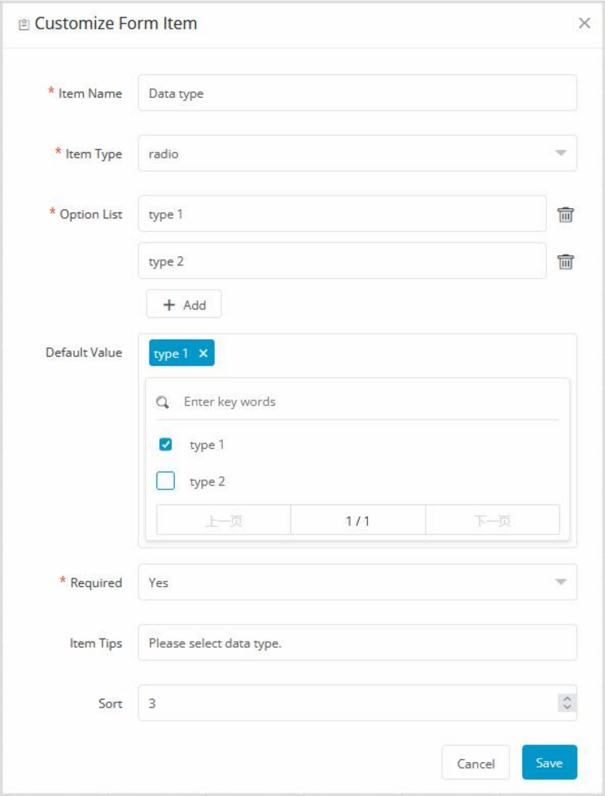

Figure 15. Customization of a radio type form item.

Item type 4: checkbox

As shown in Figure 16, the settings of a checkbox are identical to those of a radio button, but because the former is multi-choice, there is an additional max count parameter that is used to specify the maximum number of items that may be selected. This parameter is needed; if you do not wish to limit the maximum number of choices, set it to 0.

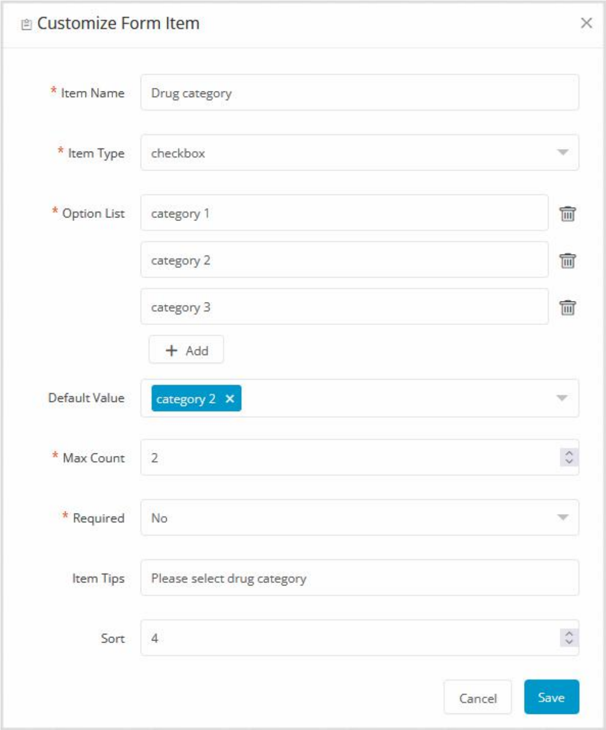

Figure 16. Customization of a checkbox type form item.

Item type 5: select

The parameters for a select type form item are identical to those for radio buttons, as illustrated in Figure 17.

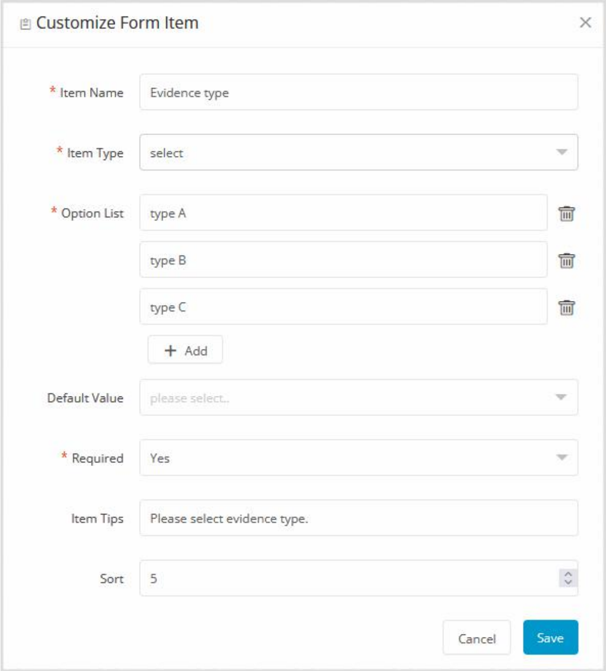

Figure 17. Customization of a select type form item.

Item type 6: xmselect

xmselect (xm-select, https://maplemei.gitee.io/xm-select/) is a multi-choice solution built on LayUI (https://github.com/sentsin/layui), a free open source web UI component library. Xmselect is a multi-selection form item, however it contains an extra option type parameter when compared to the checkbox, which is also a multi-selection type. Customize selections are available for all other option types, including checkboxes, and the xmselect option can also select a pre-set entity list, such as CancerList, DrugList, GeneList, and AlterationList (Figure 18). When these entity lists are selected, entries from the associated lists are read and the user can choose standardized entity names from them (Figure 19).

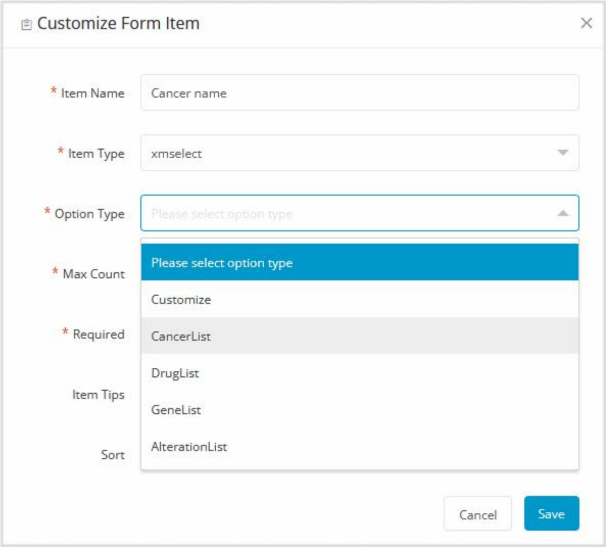

Figure 18. Option type for xmselect type form item.

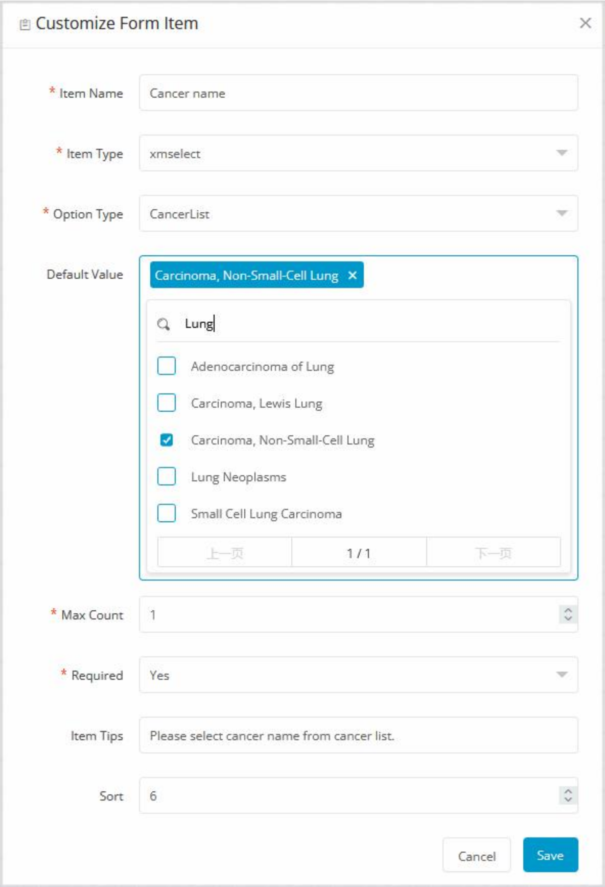

Figure 19. Customization of an xmselect type form item.

Following the creation and submission of all form items, you may preview these form items in the form details section (as shown in Figure 20). When the form is not locked, these form items can be deleted or modified, and new form items can be added. When the form is locked, all form items may only be previewed; no additions, deletions, or modifications can be made (as shown in Figure 21).

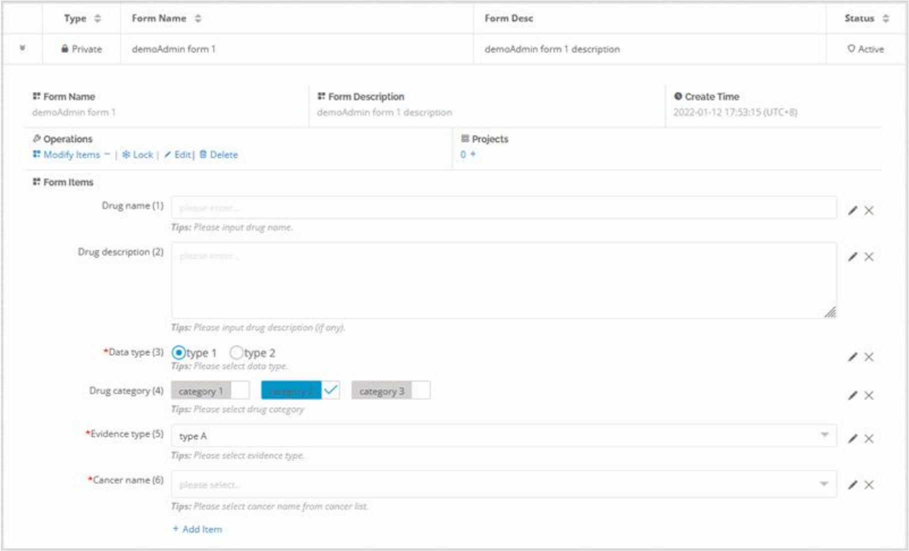

Figure 20. A form that has not been locked but has had form items generated.

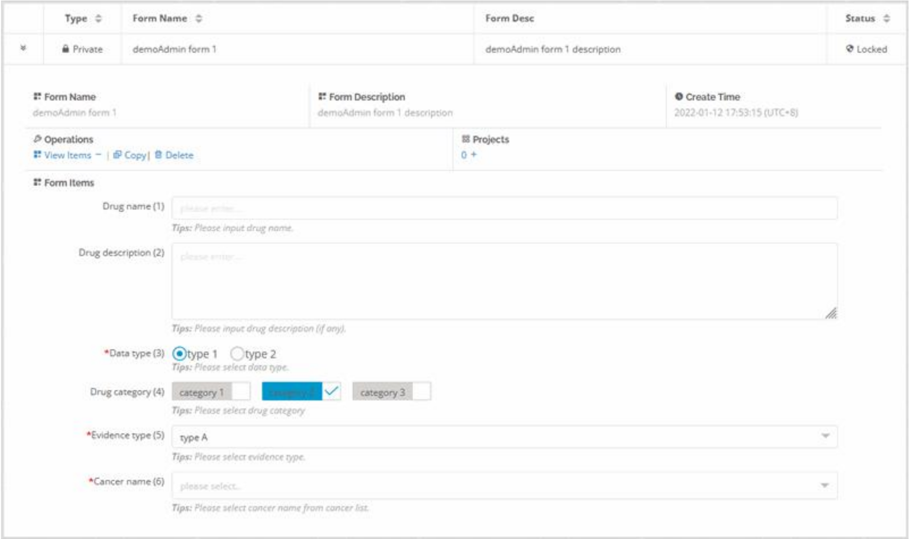

Figure 21. A form that has been locked and form items that have been generated.

It is worth noting that OncoPubMiner facilitates the collection of specific types of default settings for article data extracts, which frequently need the inclusion of the article's meta information.

For text and textarea form items, for example, you may define default values such as '[PMID]' and '[TITLE]' (as shown in Figure 22). When data is collected on the data collection page, the essential meta information may be automatically extracted (as illustrated in Figure 23), considerably reducing the labor of curators.

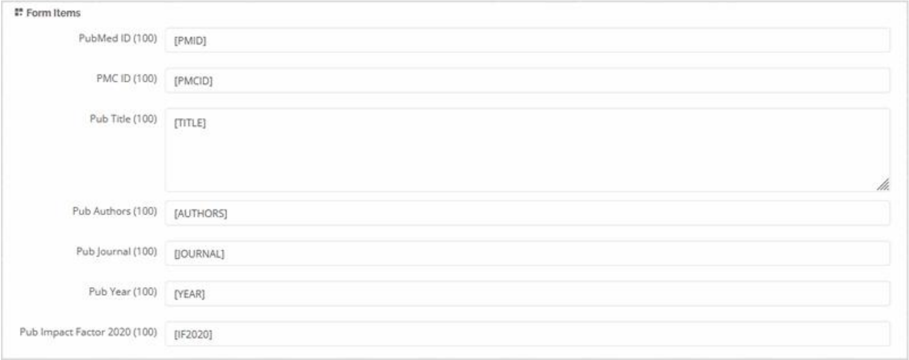

Figure 22. Default parameter settings for automatic article metadata retrieval.

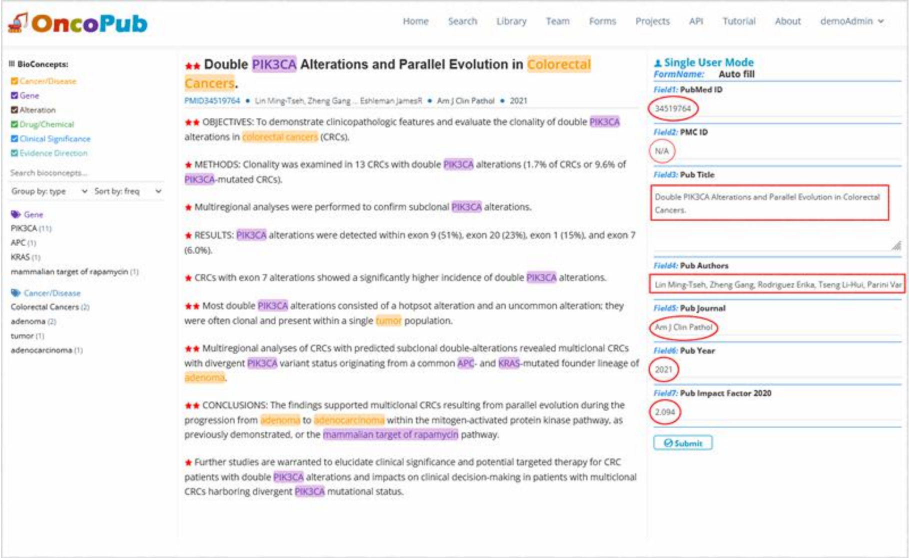

Figure 23. Metadata is taken automatically from the article data extraction page. Metadata that contains no information is padded with 'N/A'.

### 4. Projects Management

The article data mining of OncoPubMiner is primarily focused on the curation project. The project will be related with the previously constructed collection, included literature, established data structure, newly created team member accounts, and ultimately extracted knowledge data.

When the project is being established, it is required to associate the article collection and data collecting form, as well as to designate the team members who will be working on the project (as shown in Figure 24). It should be noted that only one article collection and data form may be related, whereas multiple members, all of which are set in the form of options, can be provided. It should be noted that the alternative data forms and article collections in this section can only be locked, and the unlocked ones might be modified at any moment, so they cannot be selected.

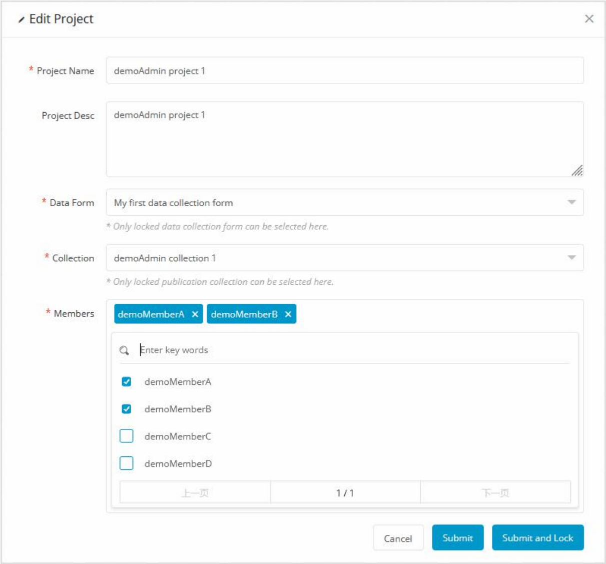

Figure 24. Create a new curation project.

After filling out or selecting the information, the user may submit and lock the project by clicking the ‘Submit and Lock’ button. Of course, you may alternatively click ‘Submit’ to complete the project's creation first, allowing you to change the details at any moment later.

Figure 25 depicts a finalized and locked project. The project's general information, as well as its associated collections, forms, and members, may be found here. Because the collection is related to literature, you may explore a list of articles that are tangentially related to the project by clicking on the link on the number below ‘Publications’. Although a project may only be attached with one form and one collection, the number of publication is infinite. Users may search for literature and add them to the collection at any moment to be connected with the project.

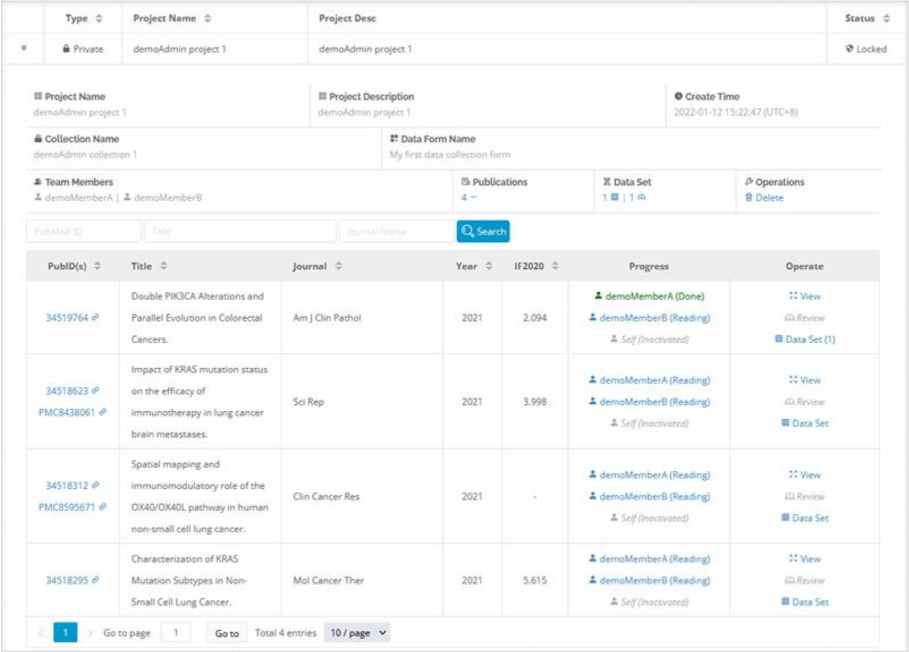

Figure 25. Overview of a created and locked project from a GA's perspective.

It is a project under the GA account, as illustrated in Figure 25, that may be utilized for curation activities at any time. Individual project members may log in to the system, open the publication list for the current project on the current page, and begin their curation job (as shown in Figure 26).

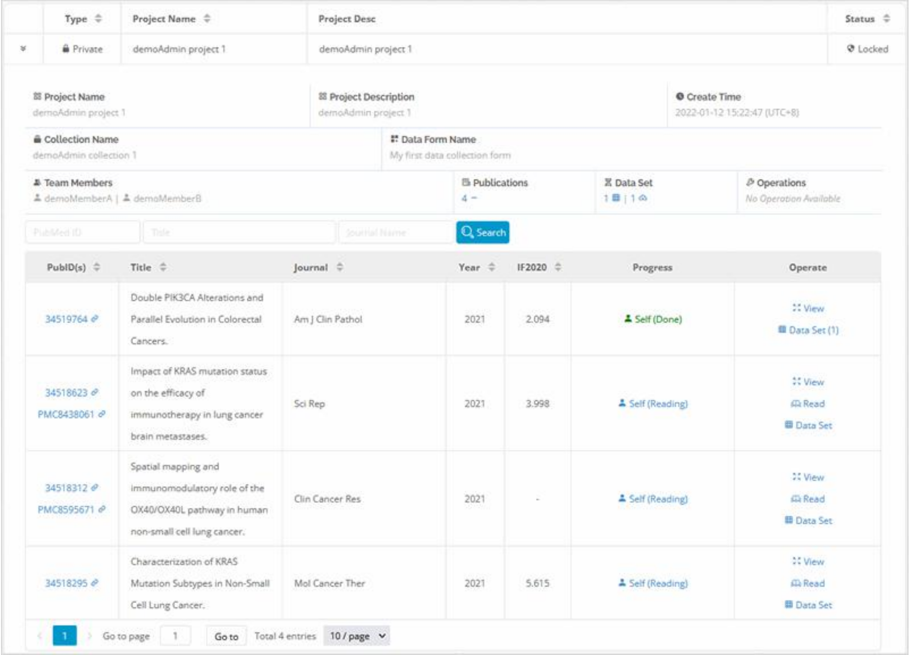

Figure 26. Overview of a created and locked project from a team member’s perspective.

When an article is not finished, the curator may see the 'Read' link behind the article, click the link to visit the data excerpt page for article reading and data gathering, and the progress field status is likewise ‘Self (Reading)’. When the article has completed extracting data, the ‘Read’ link is removed and the status is updated to ‘Self (Done)’.

It should be noted that when all project participants have finished data submission for the same article, the GA can activate the ‘Review’ link and click to reach the literature extract page for data review. The article in the project is regarded to have completed its mission only after the GA completes the data submission and clicks to get it done.

### 5. Publication Search

One of the essential and fundamental features of the OncoPubMiner system is article retrieval. This function is available to all users, whether they are signed in or out. The system's retrieval function is separated into two modes: local retrieval and distant retrieval. The former is based on local text mining findings, while the latter is based on the retrieval function given by the NCBI Eutilies API.

Remote retrieval is simple, it just calls the PubMed API, which is equal to directly accessing the PubMed database in this system. As a result, its keyword construction is identical to that of PubMed (Figure 27). After retrieving the article's PubMed ID (PMID) via the API, the system acquires the tagged BioC data in the background, adds it to the same JSON format data, and provides it to the system's front-end page for rendering and display.

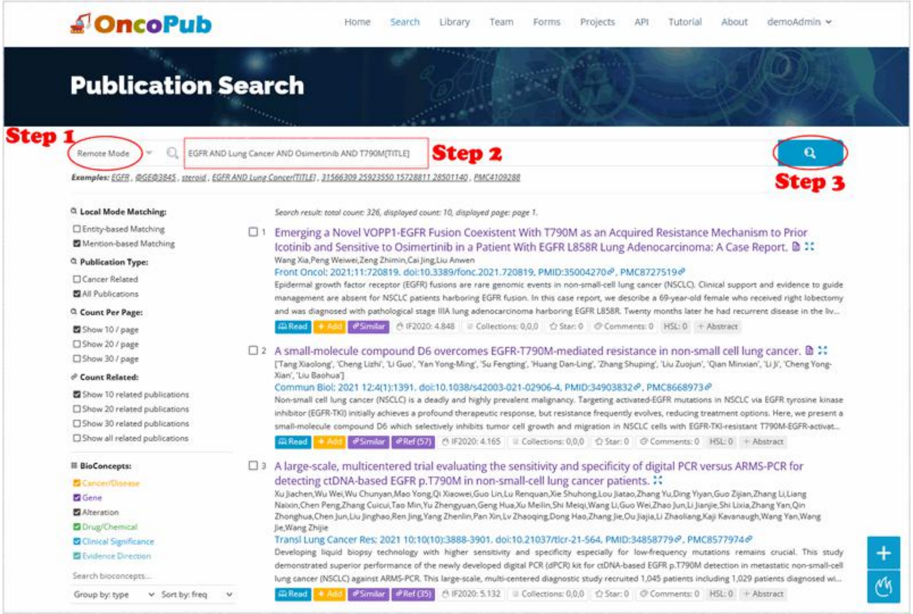

Figure 27. Remote mode-based publication search.

Based on the entity recognition processing result, the local retrieval mode is further separated into two matching methods: mention-based matching and entity-based matching. The first is a simple text matching method. When a match is found, the system background executes fuzzy matching between the string entered by the user and the entity string recognized in the article, and returns (as shown in Figure 28).

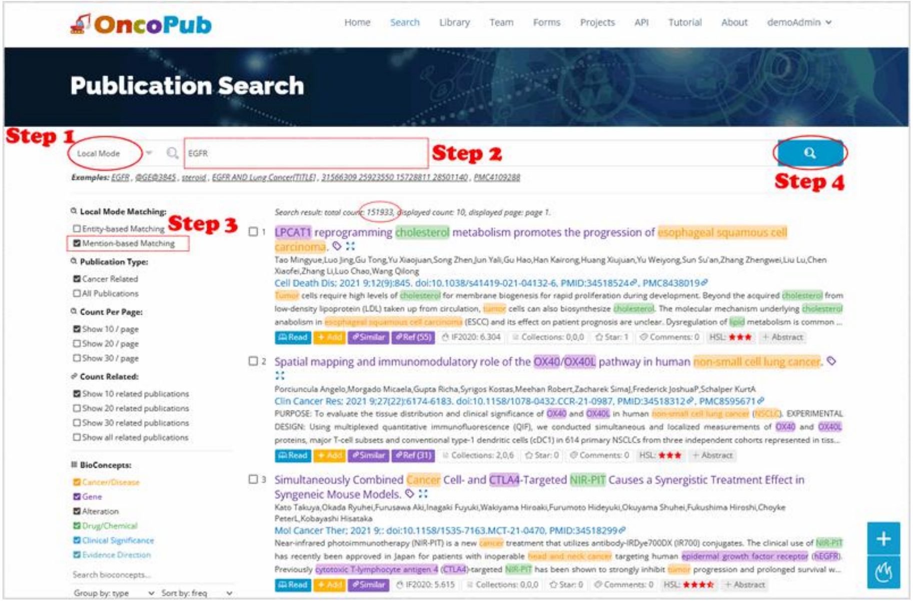

Figure 28. Local mode mention-based matching method.

The entity-based matching of local retrieval mode is a semantic retrieval strategy.

The retrieval procedure is divided into two stages. The first stage is to retrieve standard entities using keywords (Figure 29), and the second is to get proper articles using specified standard entities (Figure 30).

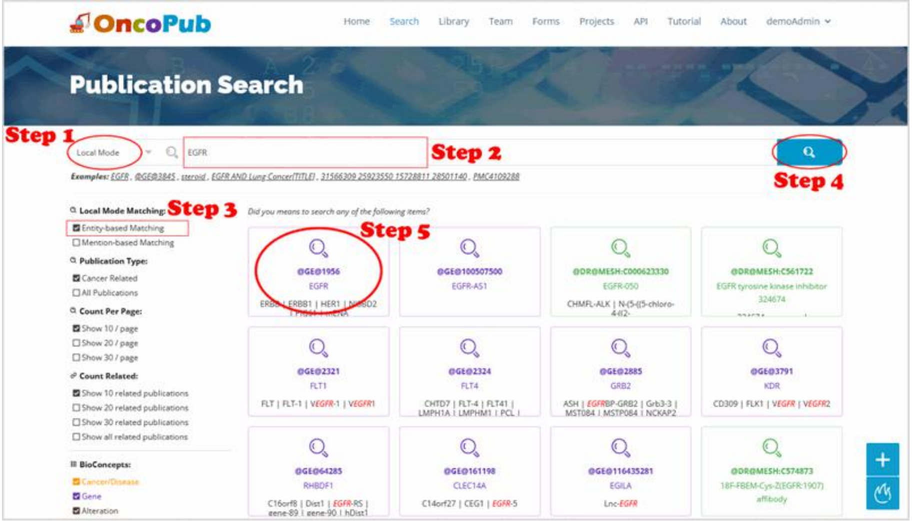

Figure 29. Local mode entity-based matching method.

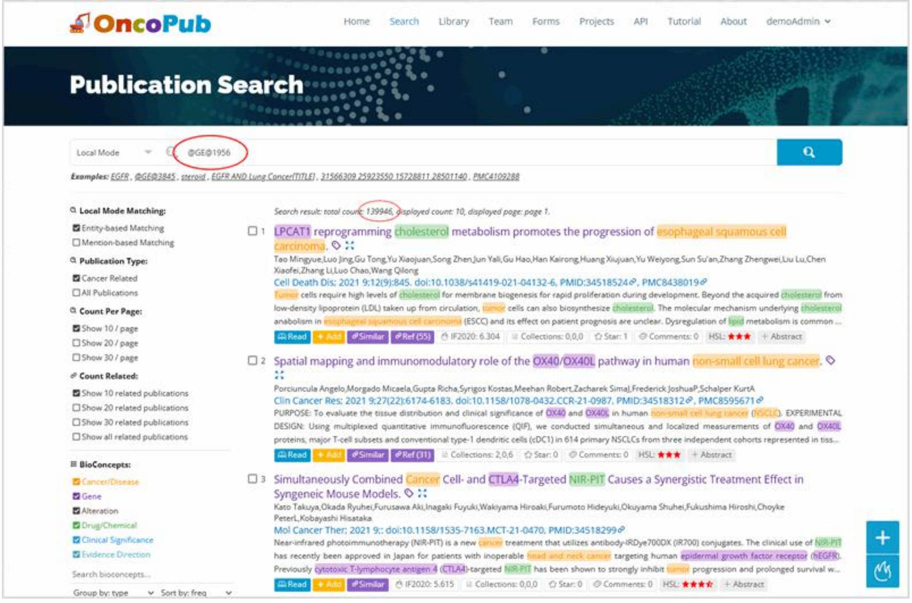

Figure 30. Use standard ID for precise retrieval based on semantic matching.

### 6. Data Collection

#### 6.1 Single User Mode

Single-user mode is a data extraction strategy that does not require an account and may be used when logged in or unlogged in. After retrieving the literature from the search page, the user clicks the ‘Read’ button and chooses the data collection form from the pop-up box to enter the data collection page for data collection (Figure 31).

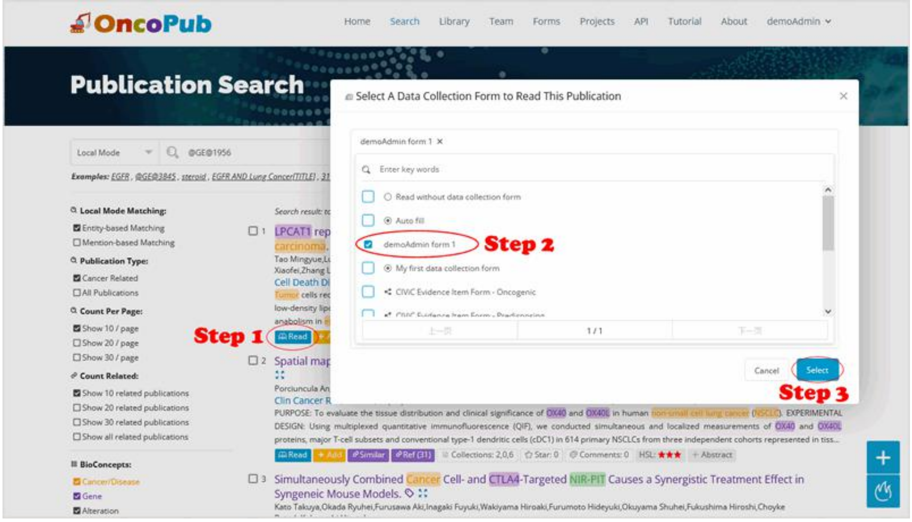

Figure 31. Begin the article reading procedure as a single user.

The user enters the article reading page, reads the article, pulls the necessary data from it, and fills out the data collecting form (Figure 32). Multiple sets of data can be submitted in batches from an article. If data has previously been submitted, it will be shown underneath the corresponding form item for user inspection. After entering all form item data and ensuring that everything is appropriate, click the submit button to submit. After the data is submitted, a page similar to the one shown in Figure 33 will appear, presenting all of the data supplied by the current user for the article, and the user can download the data by clicking the download link on the page (as shown in Figure 34).

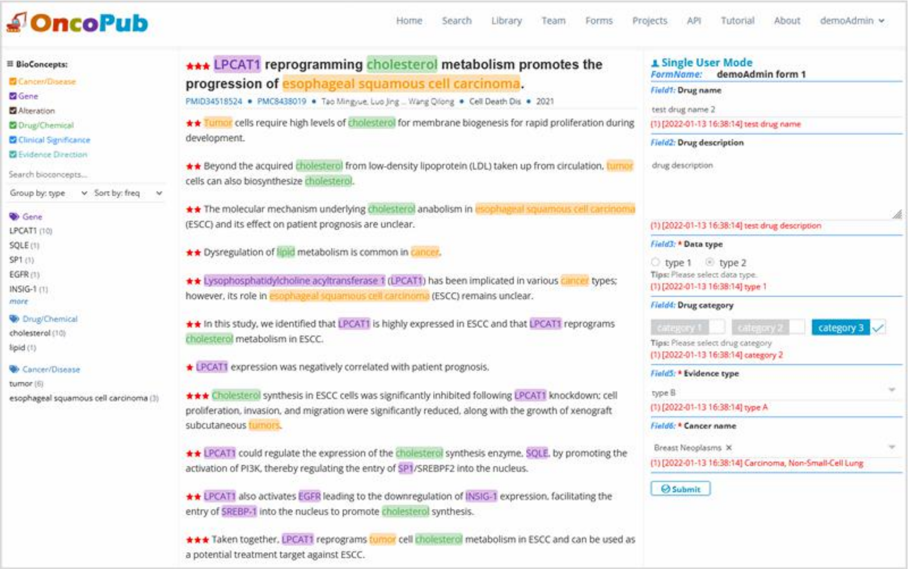

Figure 32. Sing-user mode data curation.

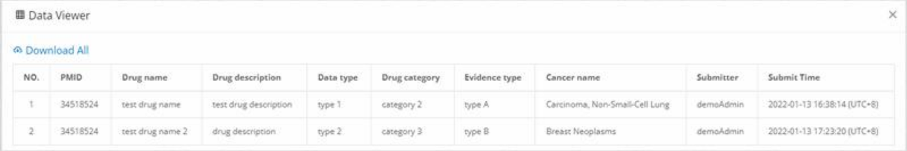
Data Viewer

Download All

| NO. | PMID | Drug name | Drug description | Data type | Drug category | Evidence type | Cancer name | Submitter | Submit Time |
|---|---|---|---|---|---|---|---|---|---|
| 1 | 34518524 | test drug name | test drug description | type 1 | category 2 | type A | Carcinoma, Non-Small-Cell Lung | demoAdmin | 2022-01-13 16:38:14 (UTC+8) |
| 2 | 34518524 | test drug name 2 | drug description | type 2 | category 3 | type B | Breast Neoplasms | demoAdmin | 2022-01-13 17:23:20 (UTC+8) |

Figure 33. Online viewer for submitted data.

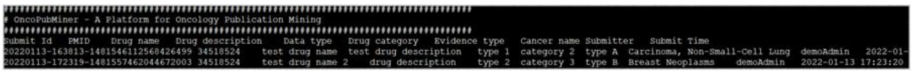

```
# OncoPubMiner - A Platform for Oncology Publication Mining
Submit Id   PMID   Drug name   Drug description   Data type   Drug category   Evidence type   Cancer name Submitter   Submit Time
20220113-163813-1481546112568426499 34518524   test drug name  test drug description   type 1  category 2  type A  Carcinoma, Non-Small-Cell Lung  demoAdmin   2022-01-
20220113-172319-1481557462044672003 34518524   test drug name 2   drug description   type 2  category 3  type B  Breast Neoplasms   demoAdmin   2022-01-13 17:23:20
```

Figure 34. Downloaded file for submitted data.

#### 6.2 Team Collaborative Mode

The team collaboration mode is also known as the project-centric mode, because all literature reading must be started from the project (as shown in Figures 25 and 26). As stated in the previous chapter on project management, the GA can only evaluate the same article once team members have read and collected data individually. The data collecting screen is presented in Figure 35 after the team members have completed the data submission and the GA has also submitted a set of data depending on the team members' submission results. Members A and B have contributed two and one pieces of data, respectively, as shown in the figure, and the time of their submission is also shown in front of the data. A data set was also supplied by the GA. This set of data can be erased and resubmitted because it was submitted by the GA himself.

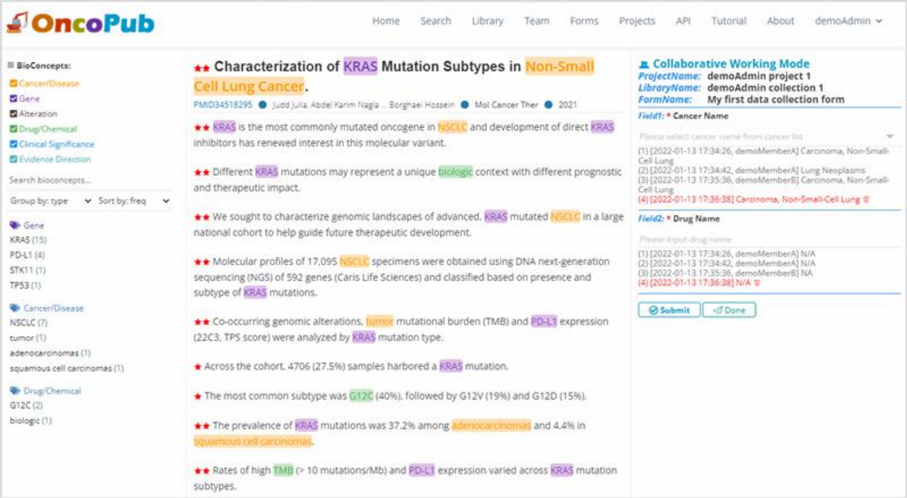

Figure 35. GA reviews the article data.

When the GA has reviewed the data and confirmed that there are no issues, he can declare the current article as complete by clicking the ‘Done’ option. Because the same article is be extracted by numerous persons and the data will be checked and submitted by the GA, the quality of the data will be considerably improved, which is critical for establishing a high-quality knowledge base.

*Last updated on 2022-02-23 11:14 (UTC+8).*
